# Loss of *UBE3A* impacts both neuronal and non-neuronal cells in human cerebral organoids

**DOI:** 10.1101/2024.02.09.579672

**Authors:** R. Chris Estridge, Z. Begum Yagci, Dilara Sen, Travis S. Ptacek, Jeremy M. Simon, Albert J. Keung

## Abstract

Angelman syndrome is a neurodevelopmental disorder caused by (epi)genetic lesions of maternal *UBE3A*. Research has focused largely on the role of *UBE3A* in neurons due to its imprinting in that cell type. Yet, evidence suggests there may be broader neurodevelopmental impacts of *UBE3A* dysregulation. Human cerebral organoids might reveal these understudied aspects of *UBE3A* as they recapitulate diverse cell types of the developing human brain. We performed scRNAseq on organoids to reveal the effects of *UBE3A* disruption on cell type-specific compositions and transcriptomic alterations. In the absence of *UBE3A*, progenitor proliferation and structures were disrupted while organoid composition shifted away from proliferative cell types. We observed impacts on non-neuronal cells, including choroid plexus enrichment. Furthermore, EMX1+ cortical progenitors were negatively impacted, disrupting corticogenesis, and potentially delaying excitatory neuron maturation. This work reveals novel impacts of *UBE3A* on understudied cell types and related neurodevelopmental processes and elucidates potential new therapeutic targets.

**Teaser:** Human cerebral organoids exhibit compositional and transcriptomic alterations in both neuronal and non-neuronal cells in the absence of *UBE3A*.

## Introduction

Human *UBE3A* is an imprinted gene located on chromosome 15^1^. While both alleles are expressed in most tissues, only the maternal allele is active in neurons due to epigenetic silencing of the paternal allele^2–4^. Loss of functional maternal *UBE3A* therefore results in a lack of UBE3A protein in neurons; this is the common lesion driving Angelman Syndrome (AS), a neurodevelopmental disorder characterized by intellectual disability, speech impairment, ataxia, and seizures^5^. Furthermore, its overexpression has been linked to other neurodevelopmental disorders including Dup15q and Autism Spectrum Disorder^6^. Thus, the dosage of *UBE3A* in the brain is likely critical for proper brain development and function^7^.

Deletion, mutation, or aberrant silencing of *UBE3A* in neurons have been associated with defects in GABAergic circuit development^8,9^, synaptic formation and plasticity^10,11^, altered neuronal proteasome^8,12–16^, and defects in dendrite^17–19^ and axon^20^ growth. This prior research focus on neurons is for three primary reasons: 1) *UBE3A* is imprinted in neurons, and thus an affected maternal allele results in almost complete loss of UBE3A protein in neurons; 2) while loss of one allele could still result in lower *UBE3A* dosage in non-neuronal cell types, with functional consequences, these cell types can be challenging to access and study, particularly those present in prenatal neurodevelopment such as neuroepithelial stem cells and progenitors; and 3) the spatial and temporal scales of neurodevelopment differ dramatically between rodent and human. Thus, while clearly important in neurons, it is important to also consider the role of *UBE3A* in other cell types, technology permitting.

Indeed, experimental and clinical evidence supports the hypothesis that there may be broader impacts of UBE3A dysregulation. In addition to its ubiquitin ligase activity, UBE3A plays a role in transcriptional regulation^21,22^ and has been reported to have centrosomal function^23,24^. Furthermore, studies showing partial paternal imprinting in neural progenitors^25^ and aberrant nuclear UBE3A in neural precursor cells^26^ suggest that the effects of altered *UBE3A* dosage may not be limited to neurons. Relatedly, conditional deletion and expression of *UBE3A* in mouse models at pre and postnatal time points indicate *UBE3A* dosage is important not only in adulthood but also during prenatal neurodevelopment^27,28^. There are also intriguing potential impacts of *UBE3A* dosage on cerebrospinal fluid (CSF), which is known to regulate both developmental and homeostatic processes of the brain^29,30^. Increased extra-axial CSF production has been observed in individuals with autism spectrum disorder^29^, and UBE3A has been found in CSF^30^ suggesting UBE3A may affect the choroid plexus (ChP), the brain region responsible for CSF production.

Human cerebral organoids could be a useful and timely model to investigate these understudied aspects of UBE3A biology as they contain the cell types that have been previously challenging to study, provide access to human molecular and cell biology, are genetically and pharmacologically perturbable, and mimic early prenatal neurodevelopmental processes and states^31–33^ including the proliferation and differentiation of stem cells into diverse neural cell types^26,31,34,35^. Our lab has previously shown these organoids can recapitulate the spatiotemporal localization and paternal silencing of UBE3A^26^. Electrophysiological studies have also revealed a potential mechanism contributing to epileptic phenotypes in AS^35^. Here we exploit organoids and single-cell RNA-sequencing (scRNAseq) to reveal the compositional and cell type-specific transcriptomic effects^32,36^ of complete and maternal *UBE3A* loss, with potential implications for understudied impacts of UBE3A on neurodevelopment, non-neuronal cells, and CSF biology. We make two primary observations. In the absence of *UBE3A*, stem cells exit multipotency earlier towards neuronal and ChP fates. In addition, neurons derived from cell lines containing partial or complete disruption of *UBE3A* exhibit disrupted corticogenesis.

## Results

### Absence of UBE3A alters the cell type composition of human cerebral organoids

Our central hypothesis is that UBE3A may have impacts on neurodevelopment beyond neuronal function. We therefore began by performing scRNAseq on whole brain human cerebral organoids derived from isogenic H9*_WT_* and H9*_UBE3A m-/p-_* pluripotent stem cells to assess how the absence of UBE3A affects cell type compositions and gene expression (Figure 1A). The H9*_UBE3A m-/p-_* cell line was generated by the Chamberlain group^37^ and contains homozygous deletions from the start site of isoform 1 to just upstream of the stop codon (exons 4-13 of transcript *UBE3A*-226, GRCh38). We confirmed the absence of *UBE3A* in H9*_UBE3A m-/p-_* organoids (Figure S1A). We chose a relatively early time point of 6 weeks to capture a variety of cell types, from stem cells to immature neurons^26^. Using 7498 high quality cells (median UMI count: 8278, median gene count: 3558; H9*_WT_*: 3376 cells, H9*_UBE3A m-/p-_*: 4122 cells), we identified 34 discrete comprising of 15 cell types consisting of mainly radial glia (RG), intermediate progenitors (IPs), mesenchymal cells (Mes), choroid plexus (ChP), RSPO+ cells, and neurons (Figures 1B, 1C, S1B, and S1C). There were RG and IP cells that also expressed cell cycle marker genes and were assigned as proliferating RG or IP. Neurons were partitioned into excitatory-like (EN) and inhibitory-like (IN) neuron clusters depending on the expression of excitatory and inhibitory marker genes (Figure S1C).

**Fig. 1.**
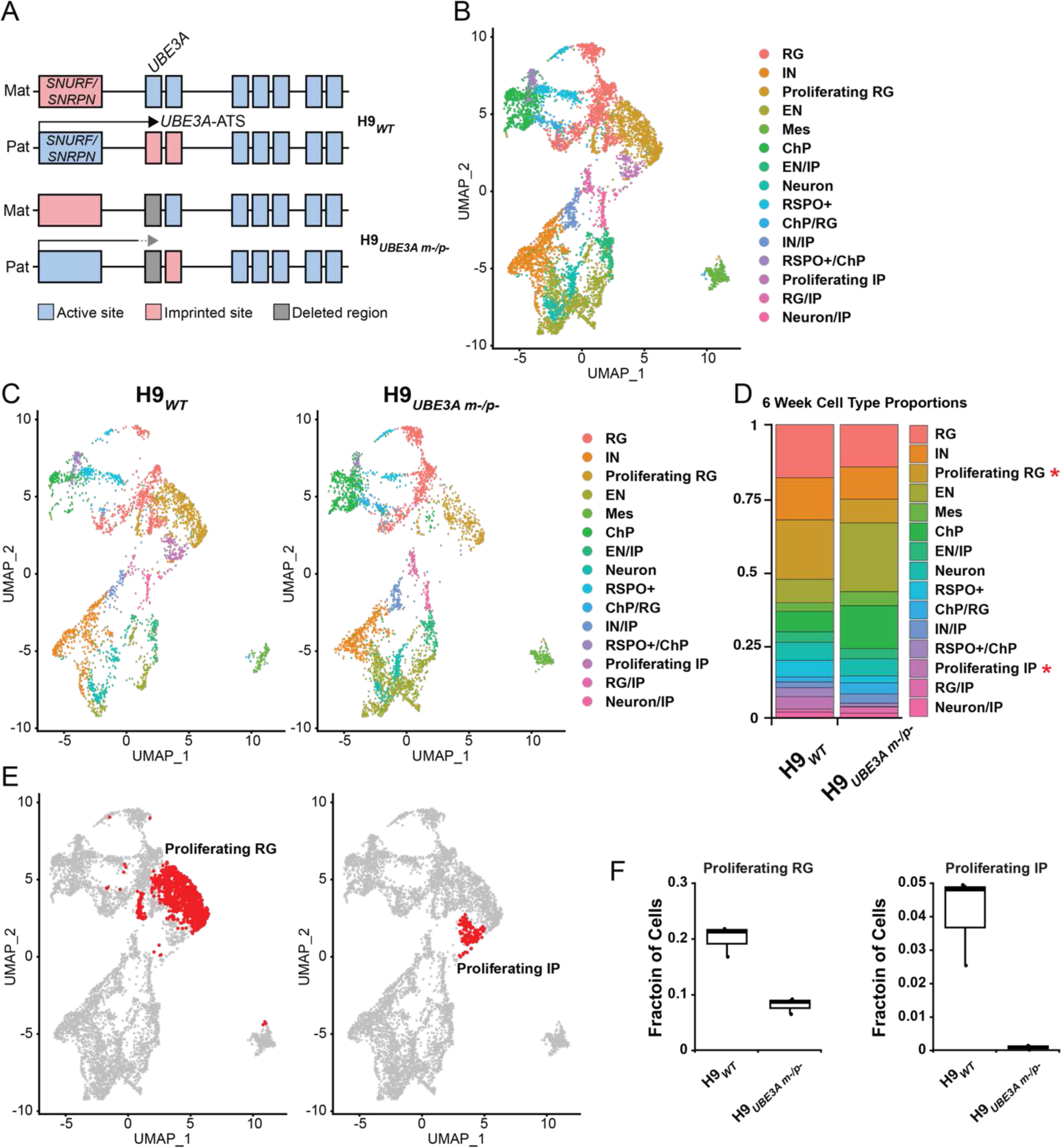
Absence of *UBE3A* alters the cell type composition of human cerebral organoids. (A) Schematics illustrating the deletion status of *UBE3A* for H9*_WT_* and H9*_UBE3A m-/p-_* cell lines used to generate organoids. (B) Uniform Manifold Approximation and Projection (UMAP) bidimensional reduction of 6 week organoid scRNAseq data with cell type identities. RG: radial glia; IN: inhibitory-like neuron; EN: excitatory-like neuron; Mes: mesenchymal; ChP: choroid plexus; IP: intermediate progenitor. (C) UMAP of 6 week H9*_WT_* and H9*_UBE3A m-/p-_* organoids. (D) Cell type proportions of 6 week H9*_WT_* and H9*_UBE3A m-/p-_* organoids. * q-value <0.05, via two-tailed t-test, n = 3 replicates of 4-5 pooled organoids per genotype. (E) UMAP with highlighting of cell types with altered proportions between H9*_WT_* and H9*_UBE3A m-/p-_* organoids. (F) Organoid composition box plots for highlighted cell types. n = 3 replicates of 4-5 pooled organoids per genotype.

Both H9*_WT_* and H9*_UBE3A m-/p-_* organoids contained each cell type. However, quantifying the proportions of cell types revealed differences in organoid compositions (Figures 1D-1F). H9*_WT_* organoids exhibited greater proportions of less mature, proliferative cell types (proliferating RG and proliferating IP) compared to H9*_UBE3A m-/p-_* organoids. Interestingly, H9*_UBE3A m-/p-_* organoids contained proportionally more mature cells like ENs, but this increase was not statistically significant. These compositional differences suggest the absence of *UBE3A* (i) might limit progenitor pools and (ii) may cause progenitors to prematurely exit multipotency and differentiate towards an EN lineage.

### H9_WT_ and H9_UBE3A m-/p-_ organoids exhibit cell type specific differences in gene expression

We performed differential gene expression analyses to inform how the absence of *UBE3A* might alter organoid compositions and how different cell types are affected. We first analyzed the sequencing reads in bulk (Figures 2A and 2B). In the absence of *UBE3A*, we observed 4206 differentially expressed genes (DEGs; 1973 down, 2233 up). This motivated further analysis for specific cell types, with EN, RG, and proliferating RG exhibiting the highest number of DEGs (Figure 2C). Qiagen Ingenuity Pathway Analysis revealed that the absence of *UBE3A* affected a plethora of canonical pathways (Figure 2D and Table S1). RG and proliferating RG cells exhibited upregulation for some neuronal signaling pathways (CREB signaling in neurons and opioid signaling pathway) and neuronal activity pathways (synaptic long term depression) while these pathways were downregulated for EN (Figure 2D and Table S1). Cell cycle and division-related pathways (e.g., cell cycle checkpoints, mitotic prophase, mitotic prometaphase, mitotic metaphase and anaphase, telomere maintenance and telomerase signaling) were downregulated for RG and ENs (Table S1). Apoptosis, senescence, and DNA damage-related pathways were also affected. RG exhibited decreased DNA damage-related pathways (role of BRCA1 in DNA damage response), apoptosis-related pathways (intrinsic pathway for apoptosis, Granzyme B signaling), and senescence-related pathways (oxidative stress induced senescence), whereas Granzyme A signaling was upregulated. Decreased telomerase signaling and increased senescence and intrinsic apoptosis pathways were present in proliferating RG. Senescence and ferroptosis signaling pathways were downregulated for ENs. Furthermore, many corticogenesis- and neurogenesis-related pathways including RHO GTPase cycle^38^, estrogen receptor signaling^39^, docosahexaenoic acid (DHA) signaling^40^, EPH-Ephrin signaling^41^, mTOR signaling^42^, reelin signaling in neurons^43^, intra-golgi and retrograde golgi to ER traffic^44^, and signaling by ROBO receptors^45^ were affected by the absence of *UBE3A* (Figure 2D and Table S1). Many cancer- and cardiac-related pathways were also deregulated by the absence of *UBE3A* (Table S1) which is not surprising due to UBE3A’s potential role in these pathways/mechanisms^46,47^. However, within the scope of the cerebral organoid model, we focused mainly on pathways related to neurodevelopment. Altogether, the absence of *UBE3A* in organoids influenced an extensive variety of pathways implicated in broad cellular processes ranging from corticogenesis to cell cycle in multiple cell types.

**Fig. 2.**
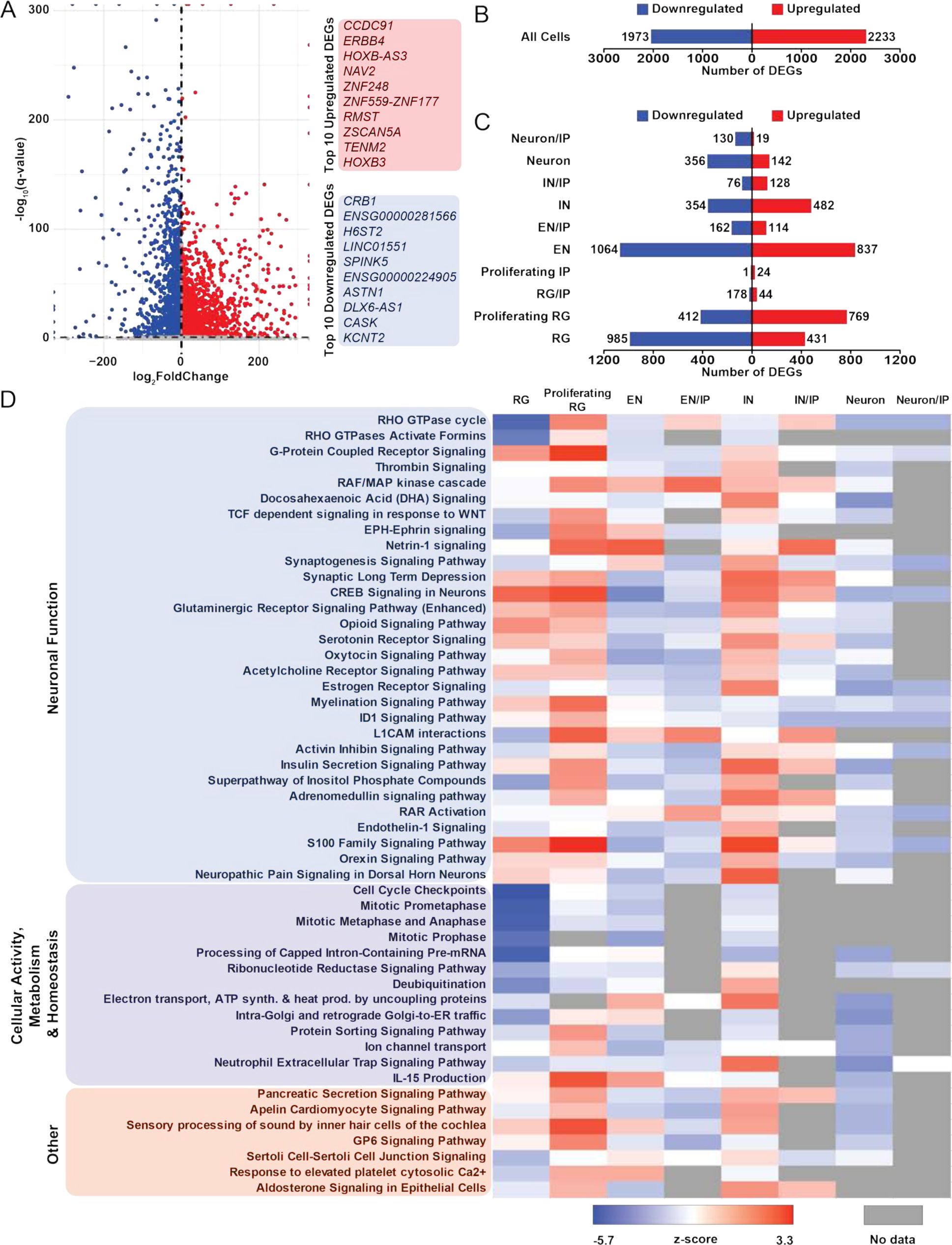
H9*_WT_* and H9*_UBE3A m-/p-_* organoids exhibit cell type specific differences in gene expression. (A) Volcano plot of bulk DEGs between H9*_UBE3A m-/p-_* and H9*_WT_* organoids. (B) Number of up and downregulated bulk DEGs between H9*_UBE3A m-/p-_* and H9*_WT_* organoids. DEG defined as log_2_FC>|1|, and false discovery rate q<0.05. (C) Number of up and downregulated DEGs separated by cell type. DEG defined as log_2_FC>|1|, and false discovery rate q<0.05. RG: radial glia; IN: inhibitory-like neuron; EN: excitatory-like neuron; ChP: choroid plexus; IP: intermediate progenitor. (D) Top 50 canonical pathways identified through QIAGEN Ingenuity Pathway Analysis (IPA) grouped by category. Differentially expressed genes with false discovery rate q < 0.05, log_2_FC) ≥|1| were used in this analysis. Pathways related to cancer or inherently from outside the nervous system were omitted. Please refer to Table S1 for all pathways.

### Absence of UBE3A disrupts progenitor proliferation and rosette structure

Our pathway analysis suggests proliferation, apoptosis and senescence might be affected (Figure 2D and Table S1). Furthermore, the absence of *UBE3A* has been linked to changes in cell proliferation and apoptosis in non-neuronal cell types^23,48,49^. Thus, as alterations in cellular processes such as cellular apoptosis, senescence, and proliferation can contribute to changes in cell type compositions, we hypothesized that the absence of *UBE3A* might drive compositional differences in organoids through one or more of these processes. Through immunostaining, we first found that there was no significant effect on apoptosis or senescence in the absence of *UBE3A* (Figure S2A). However, we did observe an impact on proliferation; we performed an EdU assay on embryoid bodies generated from H9*_WT_*, H9*_UBE3A m-/p-_* and H9*_UBE3A m-/p+_* pluripotent stem cells (Figures 3A, 3B, and S2B). The percentage of EdU-positive cells were significantly lower for H9*_UBE3A m-/p-_* and H9*_UBE3A m-/p+_* compared to H9*_WT_* organoids indicating that not only the absence of *UBE3A* but a reduced dosage of *UBE3A* leads to reduced cell proliferation in the progenitors of embryoid bodies. To check whether this inhibition of proliferation capacity happens at the pluripotent stem cell level as well, we performed an EdU assay on pluripotent stem cells (Figures 3C, 3D, and S2C). The percentage of EdU-positive pluripotent stem cells were comparable between the cell lines, suggesting that *UBE3A* impacts proliferation only of non-pluripotent stem cells and progenitors. However, interestingly, the percentage of cells in G2/M phase for H9*_UBE3A m-/p-_* pluripotent stem cells was significantly lower compared to H9*_WT_* and H9*_UBE3A m-/p+_* pluripotent stem cells, consistent with the role of UBE3A in cell cycle progression^23,49^.

**Fig. 3.**
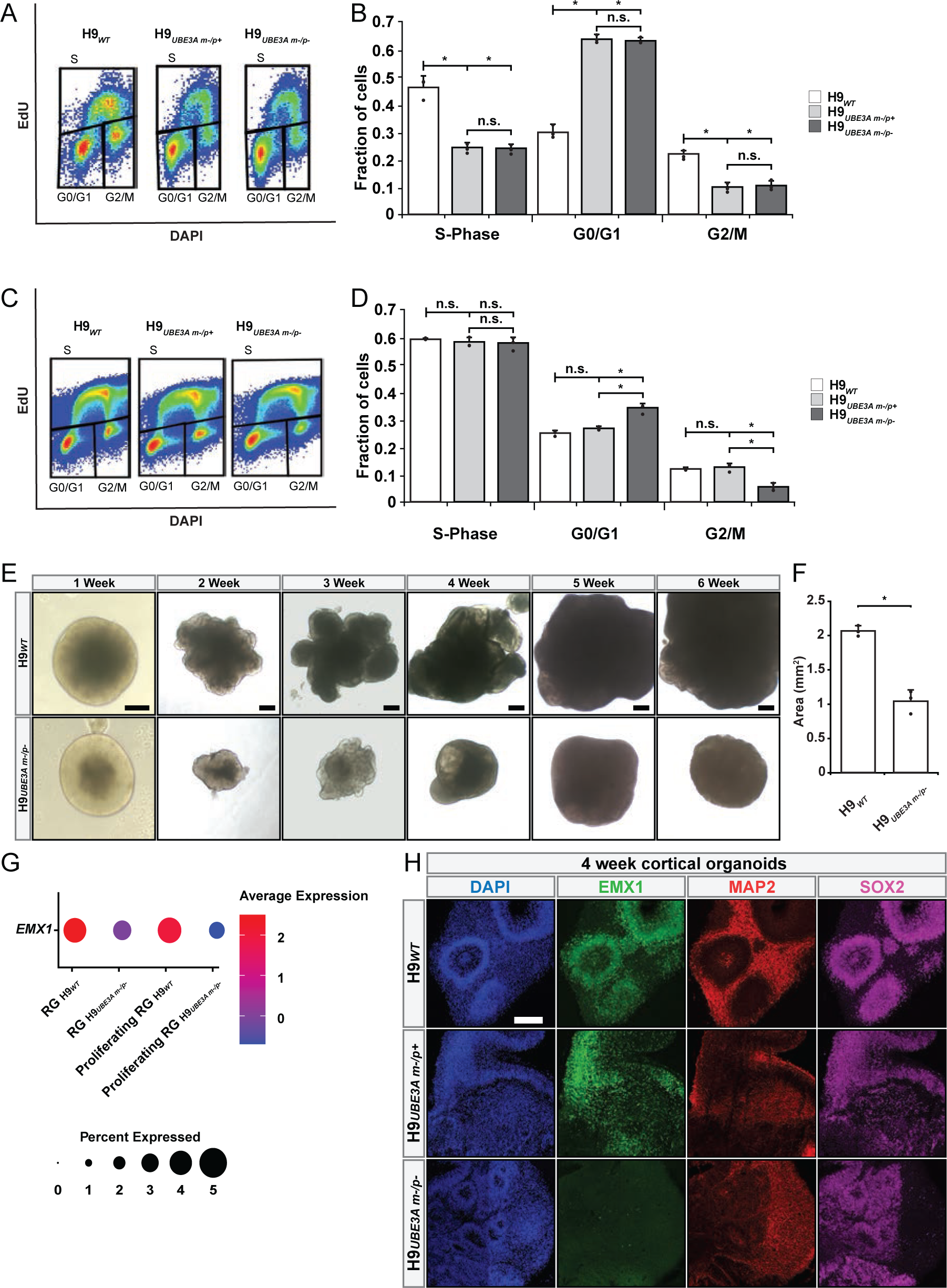
Absence of *UBE3A* disrupts progenitor proliferation and rosette structure. (A) Flow cytometry plots of EdU labeled H9*_WT_*, H9*_UBE3A m-/p+_* and H9*_UBE3A m-/p-_* embryoid bodies and (B) fraction of cells in different cell cycle states based on EdU assay performed. *p-value<0.05 via one-way ANOVA with Tukey-Kramer post hoc analysis, n.s. (not significant, p-value>0.05), n = 3 replicates of 15 pooled embryoid bodies per genotype. Full tick mark indicates sample being compared to half tick mark samples. Error bars are 95% confidence intervals. (C) Flow cytometry plots of EdU labeled H9*_WT_*, H9*_UBE3A m-/p+_* and H9*_UBE3A m-/p-_* pluripotent stem cells and (D) fraction of cells in different cell cycle states based on EdU assay. *p-value<0.05 via one-way ANOVA with Tukey-Kramer post hoc analysis, n.s. (not significant, p-value>0.05), n = 3 replicates of pluripotent stem cells from independent culture dishes. Full tick mark indicates sample being compared to half tick mark samples. Error bars are 95% confidence intervals. (E) Brightfield images of H9*_WT_* and H9*_UBE3A m-/p-_* organoid size comparison. Scale bars = 200 μm. (F) Size comparison quantification between H9*_WT_* and H9*_UBE3A m-/p-_* organoids. *p-value<0.05 via one-tailed t-test, n = 3 independent batches, each with ≥ 5 organoids per genotype. Error bars are 95% confidence intervals. (G) *EMX1* expression levels in H9*_WT_* and H9*_UBE3A m-/p-_* organoids based on 6 week scRNAseq data. (H) Immunostaining of H9*_WT_*, H9*_UBE3A m-/p+_* and H9*_UBE3A m-/p-_* human cortical organoids with EMX1, MAP2 and SOX2. Scale bars = 500 μm.

To investigate how *UBE3A* might be affecting cell proliferation, we revisited our pathway analysis. It suggested *TP53* related pathways (i.e. regulation of *TP53* activity through phosphorylation) and telomere related pathways (i.e. telomerase signaling and telomere maintenance) were altered in the absence of *UBE3A* (Table S1). UBE3A is also known to interact with p53^50^ and telomerase reverse transcriptase (TERT)^51^. Interestingly, *TP53* expression was unaffected in the H9*_UBE3A m-/p-_* organoids; however, *TERT* expression was reduced to non-detectable levels in the absence of *UBE3A* (Figure S2D). We also measured *TERT* expression levels in embryoid bodies generated from H9*_WT_*, H9*_UBE3A m-/p-_* and H9*_UBE3A m-/p+_* pluripotent stem cells. There was no significant difference between *TERT* levels in these embryoid bodies suggesting transcriptional changes occurred at a later timepoint in organoid development (Figure S2E).

We next turned our attention to potential additional downstream impacts of altered progenitor proliferation. For example, reduction in progenitor pools could result in decreased organoid size. Microcephaly is a common phenotype of AS^20,23^ and has also been observed in organoid models of other neurodevelopmental disorders^31,34,52^. We therefore measured organoid size and indeed observed H9*_UBE3A m-/p-_* organoids were significantly smaller than H9*_WT_* organoids (Figures 3E and 3F).

Neural progenitors also contribute to tissue and organ morphogenesis and structure through their architectural arrangement and generation of neurons. Our pathway analysis indicated alterations in pathways related to corticogenesis (Figure 2D and Table S1), and in particular we noticed that the cortical progenitor marker, *EMX1*^53,54^, was significantly reduced in H9*_UBE3A m-/p-_* organoids (Figure 3G). However, this dataset was generated from whole brain cerebral organoids which generate many other brain regions besides cortical tissue. To better assess the impact of EMX1 loss, which is particularly important in cortical development, we generated cortical organoids^53^ from isogenic H9*_WT_*, H9*_UBE3A m-/p-_*, and H9*_UBE3A m-/p+_* pluripotent stem cells and immunostained for EMX1 (Figure 3H). Interestingly, the dosage of UBE3A strongly impacted EMX1 levels and organization. With both *UBE3A* alleles present, EMX1^+^ progenitors organized into characteristic rosette structures (Figure 3H top). However, while H9*_UBE3A m-/p+_* organoids expressed EMX1, the organization of EMX1^+^ progenitors was severely disrupted (Figure 3H middle). Finally, in the complete absence of *UBE3A*, organoids exhibited a near total absence of EMX1 (Figure 3H bottom). While EMX1 was almost completely lost in H9*_UBE3A m-/p-_* organoids SOX2 remained (Figure 3H), indicating that the cortical progenitor population was specifically and severely affected in the absence of *UBE3A*.

*GLI3*, a radial glia marker^55^, has been reported to be upstream of *EMX* genes during dorsal telencephalon development^56^ and implicated in the sonic hedgehog (SHH) signaling pathway^56,57^. Our pathway analysis suggested downregulated SHH signaling in RG, proliferating RG, and EN cells, so we checked *GLI3* expression levels across cell types for H9*_WT_* and H9*_UBE3A m-/p-_* organoids (Figure S2F). Indeed, all cell types exhibited decreased *GLI3* expression in the absence of *UBE3A* indicating that *UBE3A* might act upstream of *EMX1* by affecting *GLI3* expression.

### Lack of UBE3A disrupts cortical layer EN development

The absence of *UBE3A* affected progenitor proliferation and organization as well as organoid size. However, we had also observed differentially expressed genes in other differentiated cell types, especially ENs, that might impact their development (Figure 2D and Table S1). To investigate these potential impacts of *UBE3A* loss, we performed scRNAseq on 11 week old H9*_WT_* and H9*_UBE3A m-/p-_* organoids as they exhibit greater proportions of differentiated and more mature cells than in the 6 week organoids we had analyzed. We chose the 11 week timepoint specifically as that is when we have previously observed a substantial increase in *UBE3A-ATS* expression^26^, the long noncoding antisense transcript that is expressed in neurons^58^ and which suppresses *UBE3A* expression^59–61^.

Using 6759 high quality cells (median UMI count: 12756, median gene count: 4558; H9*_WT_*: 4811 cells, H9*_UBE3A m-/p-_*: 1948 cells), we identified 45 discrete cell clusters comprising of 19 cell types including one unknown cell type which was excluded from further analysis (Figures 4A, S3A, and S3B). Overall, there were greater compositional differences between H9*_WT_* and H9*_UBE3A m-/p-_* organoids at 11 weeks compared to 6 weeks. In the absence of *UBE3A*, organoids contained proportionally more EN, Neuron, ChP, RG/ChP, and Mes cells and fewer Neuron/IP, EN-CL, and proliferating RG cells. Strikingly, cortical layer neurons (EN-CL) were largely absent from H9*_UBE3A m-/p-_* organoids (Figures 4B-4E, S3C, and S3D). We performed differential expression analysis to examine how the EN-CL transcriptomic profiles differed from other neurons (Figure S3E). EN-CL included many highly upregulated genes related to synapses (*SLC24A2*, *LRRC7 ANKS1B*, GRIA3/1), axon and dendrite development (*NELL2*, *NFIB*, *NEO1*), cell migration (*AFF3*, *BCL11A*), cell signaling (*PDE1A*, *GAREM1*), and highly downregulated genes related to cell differentiation (*CHD7*, *ZFHX4*) consistent with previous literature reporting the effect of *UBE3A* on these cellular functions^10,12,17,18^. H9*_UBE3A m-/p-_* organoids contained proportionally less proliferating RG consistent with the 6 week data, with the additional observation that loss of proliferating RG might lead to an increase in neurons through early exit from multipotency or a reduction in the proportion of progenitors relative to neurons. In addition, the loss of cortical layer neurons at 11 weeks appears to be an observation consistent with the loss of EMX1+ cortical progenitors at 6 weeks.

**Fig. 4.**
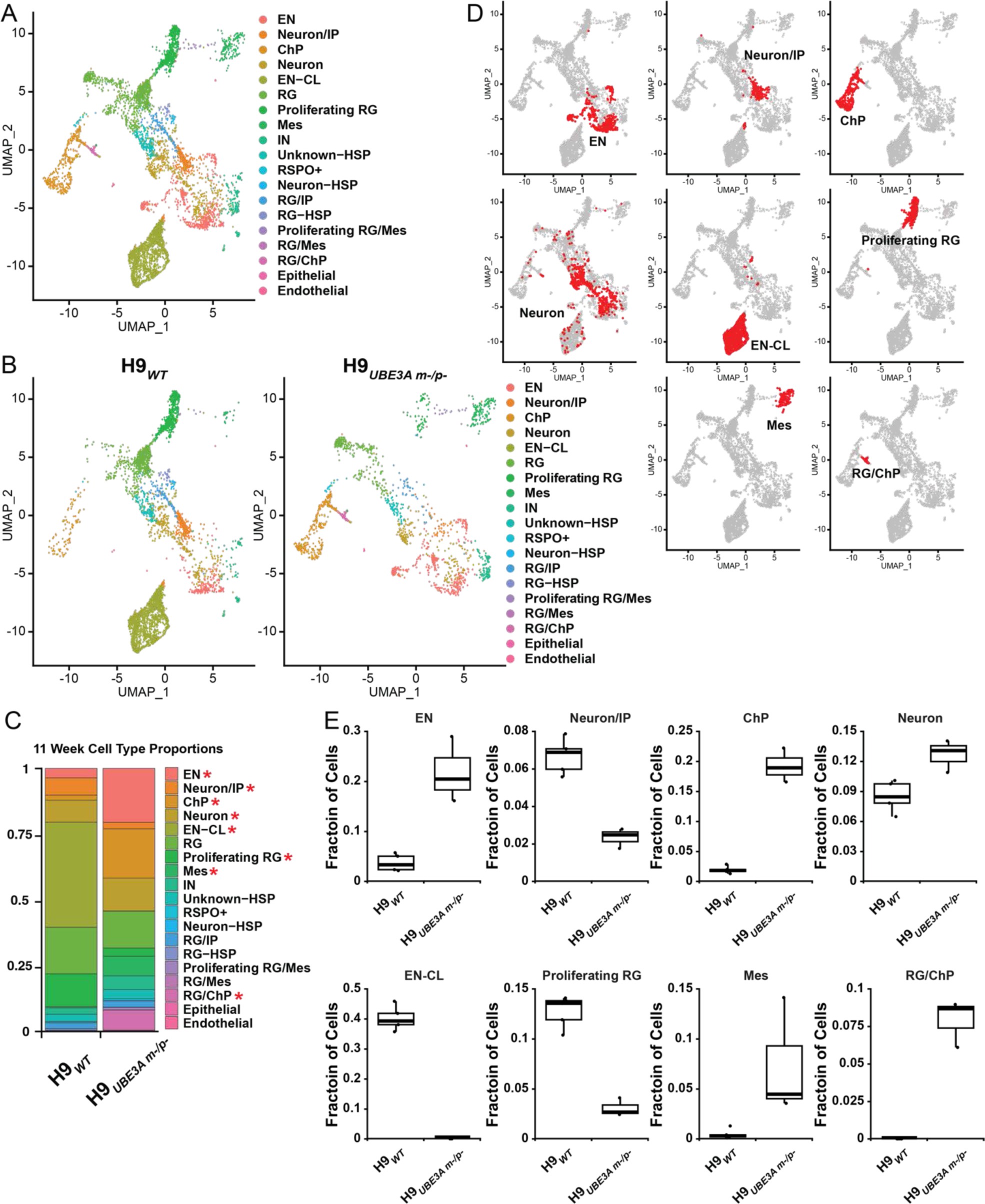
Lack of *UBE3A* disrupts cortical layer EN development. (A) UMAP bidimensional reduction of 11week organoid scRNAseq with cell type identities. RG: radial glia; IN: inhibitory-like neuron; EN: excitatory-like neuron; ChP: choroid plexus; IP: intermediate progenitor; EN-CL: cortical layer-like neurons; Mes: mesenchymal; HSP: heat shock protein. (B) UMAP of 11 week H9*_WT_* and H9*_UBE3A m-/p-_* organoids. (C) Cell type proportions of 11 week H9*_WT_* and H9*_UBE3A m-/p-_*organoids. * q-value <0.05, via two-tailed t-test, n ≥ 3 replicates of 4-5 pooled organoids per genotype. (D) UMAP highlighting cell types with altered proportions between H9*_WT_* and H9*_UBE3A m-/p-_* organoids. (E) Organoid composition box plots for highlighted cell types. n ≥ 3 replicates of 4-5 pooled organoids per genotype.

### Cell type specific transcriptomic differences between 11 week H9_WT_ and H9_UBE3A m-/p-_ ***organoids***

We performed in-depth differential gene expression and pathway analyses to investigate how the absence of *UBE3A* affected individual cell types at the 11 week timepoint. We identified 6538 DEGs across 9 cell types (Figures 5A-5C). The remaining 10 cell types did not have sufficient cell counts of each genotype for the analysis or did not contain any DEGs. Overall, more genes were downregulated than upregulated, and RG, Neuron, and ChP cells exhibited the highest number of DEGs (Figures 5B and 5C). The DEGs were used for pathway analysis. Many pathways including general signal transduction pathways (RHO GTPase cycle, EPH-Ephrin signaling, RAF/MAP kinase, ephrin receptor signaling), neuronal signaling (CREB signaling in neurons, opioid signaling), cell cycle and division (mitotic prometaphase, cell cycle checkpoints, mitotic metaphase and anaphase), circadian rhythm-related (orexin signaling pathway) and neuron-related pathways (synaptogenesis signaling pathway, myelination signaling pathway) were affected by the absence of *UBE3A*. IPA revealed that there were cell-type specific differences in these pathways (Figure 5D and Table S2). The most salient observation was that inhibitory-like neurons (IN) upregulated genes associated with several signaling pathways while most other cell types repressed these pathways. Although performed at different times and therefore subject to batch effects, there were qualitative differences between 6 week and 11 week organoids (Figures 2D, 5D and Tables S1, S2). This difference was more obvious in proliferating RG where general signal transduction pathways involving RHO GTPases, EPH-Ephrin, RAF/MAP kinase, and CREB were all downregulated in 11 week organoids but upregulated in 6 week organoids. Additionally, more cell cycle and division related pathways were downregulated in 11 week compared to 6 week organoids.

**Fig. 5.**
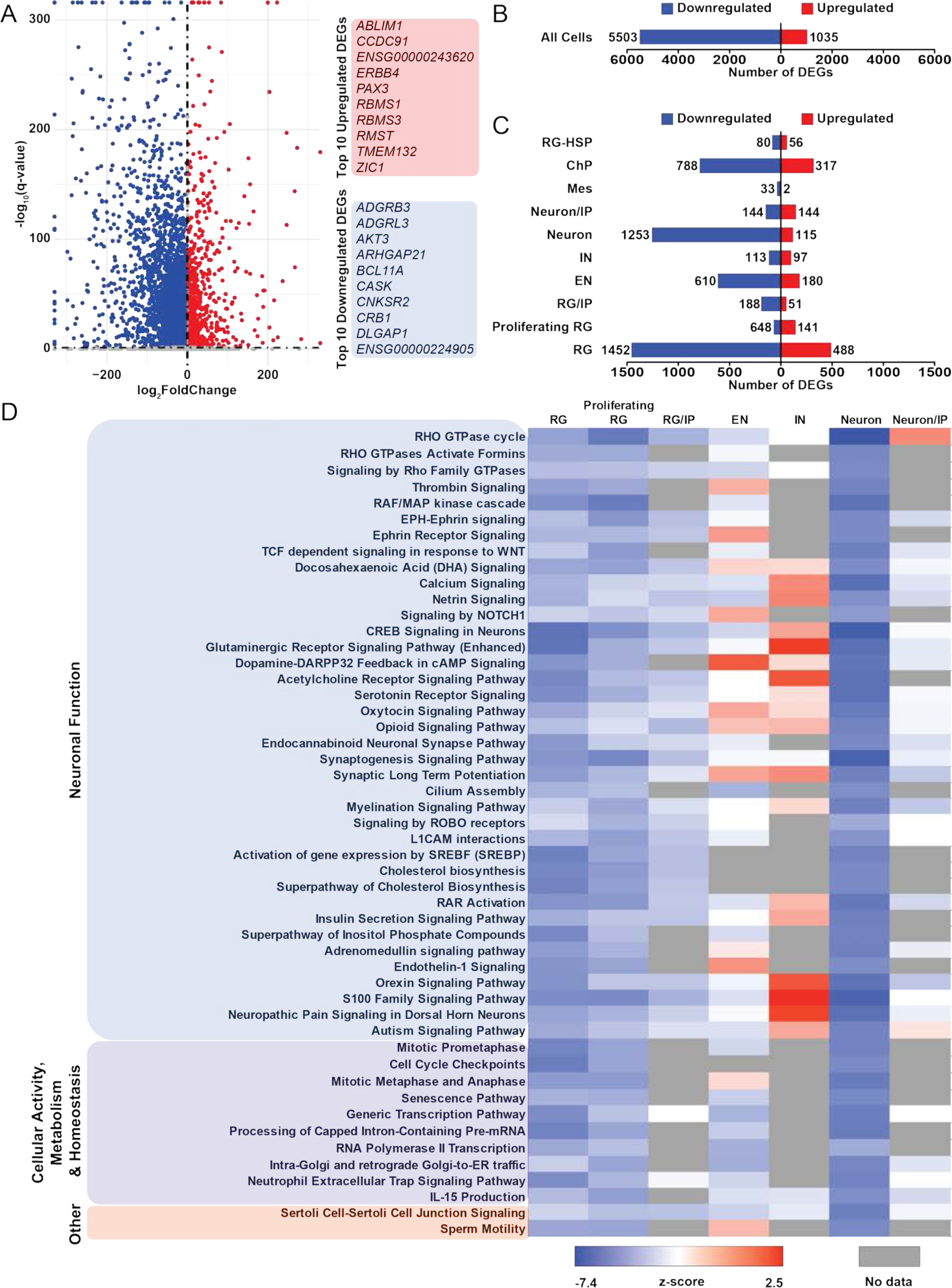
Cell type specific transcriptomic differences between 11 week H9*_WT_* and H9*_UBE3A m-/p-_* organoids. (A) Volcano plot of bulk DEGs between H9*_UBE3A m-/p-_* and H9*_WT_* organoids. (B) Number of up and downregulated bulk DEGs between H9*_UBE3A m-/p-_*and H9*_WT_* organoids. DEG defined as log_2_FC>|1|, and false discovery rate q<0.05. (C) Number of up and downregulated DEGs separated by cell type. DEG defined as log_2_FC>|1|, and false discovery rate q<0.05. RG: radial glia; IN: inhibitory-like neuron; EN: excitatory-like neuron; ChP: choroid plexus; IP: intermediate progenitor; Mes: mesenchymal; HSP: heat shock protein. (D) Top 50 canonical pathways identified through QIAGEN Ingenuity Pathway Analysis (IPA) grouped by category. Differentially expressed genes with false discovery rate q < 0.05, log_2_FC ≥|1| were used in this analysis. Pathways related to cancer or inherently from outside the nervous system were omitted. Please refer to Table S2 for all pathways.

### Excess ChP generation in the absence of UBE3A

In addition to observed impacts on progenitors and neurons, we also observed alterations in organoid composition and DEGs related to ChP cells, including an increased proportion of ChP cells in H9*_UBE3A m-/p-_* organoids and a substantial number of DEGs compared to H9*_WT_* ChP cells. The ChP comprises the brain tissue that produces CSF. Recent human clinical studies also suggest a link between UBE3A and CSF^29,30^. UBE3A has been detected in CSF^30^, and individuals with Autism Spectrum Disorder (which has high comorbidity with AS) exhibit increased extra-axial CSF production^29^. Given this set of experimental and clinical observations, we performed additional experiments to better characterize the phenotypes associated with complete loss of *UBE3A* as well as identify what phenotypes might arise in organoids generated from a clinically relevant genotype, H9*_UBE3A m-/p+_*.

We generated H9*_WT_*, H9*_UBE3A m-/p-_*, and H9*_UBE3A m-/p+_* organoids and observed two discrete morphological phenotypes through brightfield microscopy: large fluid-filled cysts^62,63^ and sinuous/invaginated tissue^64^, both characteristic of organoids with ChP tissue (Figures 6A and 6B). Immunostaining for TTR confirmed the ChP identity of the cysts and sinuous tissue (Figure 6C). Interestingly, UBE3A dosage appeared to control the balance of the two morphologies, with H9*_WT_* containing cysts, H9*_UBE3A m-/p-_* containing sinuous tissue, and H9*_UBE3A m-/p+_* containing both morphologies.

**Fig. 6.**
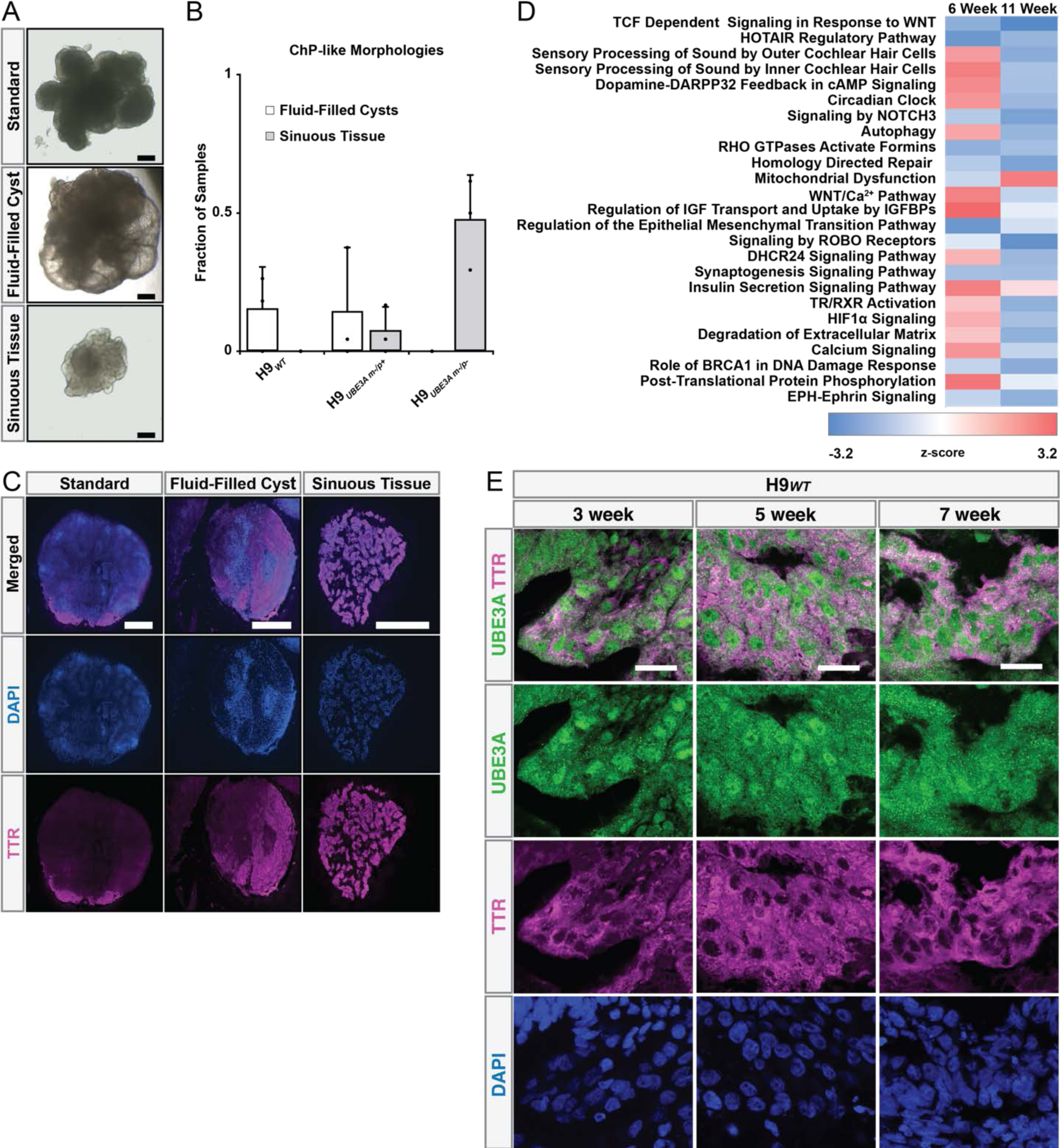
Excess ChP generation in the absence of *UBE3A*. (A) Brightfield images showing fluid-filled cyst and sinuous tissue morphologies in organoids. Scale bars = 200 μm. (B) Quantification of ChP-like morphologies. n = 3 independent organoid batches each with ≥ 5 organoids per genotype. CHP: choroid plexus. Error bars are 95% confidence intervals. (C) TTR immunostaining in fluid-filled cysts and invaginations. Scale bars = 500 μm. (D) Canonical pathways deregulated by the absence of *UBE3A* in 6 week and 11 week organoid scRNAseq data. Top 25 canonical pathways are shown. Differentially expressed genes with false discovery rate q < 0.05, log_2_FC ≥|1| were used in this analysis. Pathways related to cancer or inherently from outside the nervous system were omitted. Please refer to Tables S1 and S2 for all pathways. (E) Spatiotemporal localization of UBE3A in ChP-like tissue. Distinct nuclear UBE3A observed in early H9*_WT_* organoids weakens over time in TTR+ ChP cells. Scale bars = 25 μm.

The difference in proportion of ChP cells as well as the morphological differences of the ChP tissue suggested that loss of *UBE3A* may have functional impacts. We therefore performed pathway analysis on the scRNAseq data from both the 6 and 11 week organoids for the ChP cells (Figure 6D, Tables S1 and S2). The absence of *UBE3A* affected the expression of many WNT-related pathways (*e.g.*, TCF dependent signaling in response to WNT, HOTAIR regulatory pathway), circadian clock, autophagy and mitochondria-related pathways (*e.g.*, mitochondria dysfunction) consistent with literature^15^.

Some pathways showed differences between the 6 and 11 week time points, such as autophagy (Figure 6D). Given the dynamic nature of UBE3A expression as well as localization^26^, this motivated us to examine where and when UBE3A was present in the ChP cells. We generated 3, 5, and 7 week old H9*_WT_* organoids and immunostained for both TTR and UBE3A (Figure 6E). This revealed strong nuclear localization of UBE3A in TTR+ epithelial cells of ChP-like regions in 3 and 5 week old organoids. However, this pattern was lost in older 7 week old organoids with UBE3A becoming diffuse across subcellular compartments.

## Discussion

The primary conclusion of this work is that *UBE3A* may have multivariate and wide-ranging impacts on the developing brain, well beyond the previously considered impacts on the synaptic functions of adult neurons. More specifically, we showed the absence of *UBE3A* affected the composition, morphologies, functions, and gene expression of both neuronal and non-neuronal cell types and tissues. Here we present several models or interpretations consistent with our experimental results and their potential downstream clinical impacts that might serve as the basis for future experimental and clinical work.

First, we posit that *UBE3A* may help regulate the overall compositional balance and size of the brain through direct impacts on neural stem and progenitor cell proliferation. This may connect to observations of microcephaly in AS (albeit a phenotype with incomplete penetrance), as well as to potential imbalances in neuronal subtypes that might affect relative excitatory and inhibitory signaling. Models of other neurodevelopmental disorders have also reported altered organoid compositions^34,65^. Our results suggest *UBE3A* helps support neural stem and progenitor proliferation potentially through impacts on cell cycle progression. The absence of *UBE3A* resulted in a greater proportion of cells in G0/G1 phase indicating possible cell cycle arrest. Several studies have also reported a potential role for UBE3A in cell cycle progression in other non-neural cell types^23,49^: *UBE3A* expression is cell cycle dependent and *UBE3A* knockdown leads to defects in chromosome segregation and spindle dynamics^23^; UBE3A has also been shown to regulate the levels of cyclin-dependent kinase inhibitor p27, and reduction of UBE3A increases p27 levels and cell cycle arrest^49^. It has also been reported that UBE3A may influence apoptosis and senescence through its protein interactions with p53 and TERT^50,51^, and increased apoptosis has been observed in neural precursor cells derived from an AS mouse model and attributed to increased reactive oxygen species^66^. In the absence of *UBE3A*, we observed a transcriptional reduction of TERT expression below the detectable threshold in 6 week organoids, while we did not detect alterations in TERT levels in embryoid bodies. We were not able to detect altered rates of apoptosis or senescence and did not observe a significant transcriptional difference in p53 level between H9*_WT_* and H9*_UBE3A m-/p-_* organoids; however, we cannot discount the possibility these processes may be affected at transcriptional levels undetectable in our model system and assays or at protein level.

Second, we propose a model where *UBE3A* negatively impacts signaling through the *SHH-GLI3-EMX1* axis with a resultant disruption of corticogenesis that might be connected to altered cortical activity and structure in AS. As described above, we observed that neural stem and progenitor proliferation was impacted, but these neural precursor cells are not homogeneous. Therefore, pathway analysis led us to specifically investigate cortical precursor cells which are characterized by their expression of *EMX1* and downstream corticogenesis. Indeed, the absence of *UBE3A* resulted in a significant decrease in the *EMX1+* cell population as well as disruption of neural rosette. Providing additional confidence in this functional connection, these phenotypes were dependent on UBE3A dosage.

These observations may also be consistent with the disruption of neuronal maturation observed in other model systems, although the underlying mechanisms remain unclear in all cases^67,68^. In addition, to our knowledge, this is the first report of an association between *UBE3A* and *EMX1*. Therefore, we explored the DEGs and pathway analysis more deeply to identify what pathways might be affected upstream of *EMX1*. Outside of neural contexts, UBE3A has been associated with many pathways related to corticogenesis and neurogenesis such as mTOR^69^, Ephrin^11^, and WNT/β-catenin^70^. However, our differential gene expression analysis implicated SHH signaling. There is some mechanistic precedent for this potential connection between a ubiquitin ligase and SHH signaling: RNF220, a ubiquitin E3 ligase has been reported to modulate the SHH/GLI signaling gradient by interacting with GLI proteins through ubiquitination^57^. Interestingly, we observed that the expression levels for *GLI3*, which is upstream of *EMX* genes^56^, to be lower in the absence of *UBE3A*. This is just one possibility of how *UBE3A* participates in the regulation of corticogenesis, and future work might assess this and the other affected pathways identified in our and others work.

It will also be prudent to note that *EMX1+* progenitors were not the only class of progenitors affected by the loss of *UBE3A*. Proliferating RG in general were not only negatively impacted, but interestingly in 6 week old organoids they also upregulated neuron-related pathway genes. This might indicate that these neural precursors are proceeding through asymmetric division and exiting multipotency earlier than normal.

Third, we suggest that UBE3A is involved in ChP generation and function with potential relevance to differences in CSF volume observed in individuals with autism spectrum disorder^29^. This connection is perhaps the most tenuous to make, yet perhaps also the most intriguing. We observed increased ChP generation in the absence of *UBE3A*, as well as DEGs associated with IPA pathways previously linked to ChP including WNT-related pathways, mitochondrial dysfunction^71,72^, circadian clock^73^, and autophagy^74^. We also observed an interesting transient nuclear localization of UBE3A within ChP in organoids. Although the subcellular localization of UBE3A is likely related to its function^26^, it is unclear if this transient spatiotemporal localization pattern is functionally relevant.

*UBE3A* expression has been detected in developing and adult human ChP in various databases (Table S3). It remains unclear how UBE3A might be mechanistically linked to ChP generation and function as well as what the functional consequences are for AS individuals; however, the potential implications could be important as the ChP regulates clearance and secretion of important regulatory factors both in adults and during brain development. This might also include UBE3A protein that has been detected in the CSF^31^ and that could be distributed throughout the brain via the CSF and enter other cells^75^. It might also impact the generation, processing, and clearance of amyloid beta related to Alzheimer’s Disease pathology; previously, a study showed that UBE3A-deficient Alzheimer’s Disease model mice exhibited decreased amyloid beta levels^76^. Potentially consistent with this observation, the amyloid processing pathway was upregulated in 11 week ChP cells in organoids deficient for *UBE3A* (Table S1).

We end our discussion with two forward-looking technological notes. First, we discuss the specific utilities and limitations of the organoid models presented here. It is important to realize that the genetic perturbations and organoids used in this work are not directly reflecting physiological conditions. They cannot capture behavioral outcome measures as animal models can nor complex phenotypes arising from a complete organismal physiology. Instead, the strengths of organoids are in providing direct and clean perturbations while maintaining a high level of complexity in cell type, tissue structures, and human specific (epi)genetics. This has additional important implications. For example, not only is a double knockout of *UBE3A* embryonic lethal in mice, even the use of conditional deletions later in rodent development would be subject to many potential compensatory mechanisms present in a full organism but not in an organoid. A corollary is that while a double knockout of *UBE3A* is non-physiological, it can uncover important biology that would otherwise be hidden from view or below detectable thresholds using existing analytical technologies, or in most animal and tissue culture models of disease. In other words, with the inherent lack of compensatory mechanisms present in cerebral organoids compared to a full organism, molecular and cellular “burdens” induced by pathological lesions may be exacerbated, uncovering novel biology that could be difficult if not impossible to observe in other model systems.

A second forward-looking note is that despite these scientifically useful aspects of organoids, it will remain important to explore how these findings might translate to animal models as well as patient derived samples. For example, here we focused on isogenic pluripotent cell lines with *UBE3A* completely absent. This provided the cleanest and most direct experimental approach to assess *UBE3A* biology in organoids. However, an important next step could be to investigate the findings from this work in the context of diverse AS induced pluripotent cell lines reflecting the multiple different AS etiologies (large deletion, mutation, uniparental disomy, imprinting center defects, mosaicism). This will be important in comparing phenotypes arising in a heterozygous background where paternal *UBE3A* is still present, as well as in parsing the contributions of other genes deleted or silenced in the different (epi)genotypes of AS.

## Materials and Methods

### hESC lines

Feeder-independent H9 human embryonic stem cells (hESCs) (WA09) were originally obtained from WiCell. H9*_UBE3A m-/p-_* and H9*_UBE3A m-/p+_* cells with a 66 kb deletion (GRCh38/hg38 chr15: 25339158-25405461) were derived from WA09 and provided by Dr. Stormy Chamberlain (UCONN)^37^. Our group also confirmed the deletion in these cells in an earlier study^26^. Cells were maintained in tissue culture dishes (Fisher Scientific Corning Costar) coated with 0.5 mg/cm^2^ Vitronectin (VTN-N) (Thermo Fisher Scientific) in E8 medium (StemCell Technologies) and passaged using standard protocols. Cells were maintained in a humid incubator at 37°C with 5% CO_2_.

### Cerebral organoid generation

Whole brain and cortical human cerebral organoids were generated and maintained using the same protocols as described^31,53,77^. Organoids were maintained in a humid incubator at 37°C with 5% CO_2_.

### Cryosectioning and immunohistochemistry

Tissues were fixed in 4% paraformaldehyde for 15 min at 4°C followed by 3 x 10 minutes PBS washes. Tissues were allowed to sink in 30% sucrose overnight and then embedded in 10% gelatin/7.5% sucrose. Embedded tissues were frozen in an isopentane bath between −50 and −30°C and stored at -80°C. Frozen blocks were cryosectioned to 30 μm. For immunohistochemistry, sections were blocked and permeabilized in 0.3% Triton X-100 and 5% normal donkey serum in PBS. Sections were incubated with primary antibodies (Table S4) in 0.3% Triton X-100, 5% normal donkey serum in PBS overnight at 4°C in a humidity chamber. Sections were then incubated with secondary antibodies in 0.3% Triton X-100, 5% normal donkey serum in PBS for 2 hr at RT, and nuclei were stained with DAPI (Invitrogen). Slides were mounted using ProLong Antifade Diamond (Thermo Fisher Scientific). Secondary antibodies used were donkey Alexa Fluor 488, 546 and 647 conjugates (Invitrogen, 1:500).

Images were taken using a Nikon AR confocal laser scanning microscope (Nikon Instruments). All samples within quantification experiments were imaged using the same laser intensity settings using the 10X objective. Quantifications were performed manually in FIJI. Sample names were first removed, and the images randomized before quantification to avoid bias. A region of interest was generated around the organoid DAPI outline. The area containing both marker of interest and MAP2 within the organoid was then calculated. the cell death was calculated from the area fraction of marker of interest and MAP2 relative to total MAP2 area within the organoid. At least 3 organoids per condition were collected and measured.

### Single cell dissociation of organoids for scRNAseq

Organoids were dissociated using a previously described protocol^53,78^ which is a modification of the Worthington Papain Dissociation Kit manufacturer’s protocol (Worthington, #LK003153). Briefly, reagents were resuspended according to manufacturer’s instructions. Organoids were placed into papain and DNAse mix and minced with a sterile razor. Minced organoids were incubated for 30 min at 37°C. Following this, minced pieces were broken up by pipetting up and down with p1000 pipette and incubated for 10 min at 37°C. Next, the dissociated organoid mix was placed into inhibitor solution and passed through a 40 μm filter, centrifuged at 300 g for 7 min, and resuspended with 1x ice cold PBS and kept on ice. Cells were counted with an automated hemocytometer (Invitrogen) and single cell formation was confirmed with an inverted tissue culture microscope (Nikon) before proceeding to the fixation.

### scRNAseq library preparation, sequencing, and analysis

Fixation, cell barcoding, and library preparation was done using a previously described split-pool approach^79^ with a commercial kit (Parse Biosciences). Briefly, 3,500,000 cells were fixed per sample using the manufacturer’s instructions. Approximately 13,500 cells/well were seeded in first barcoding plate and cDNA was generated with an in-cell reverse transcription reaction using well-specific barcodes. Cells were then pooled and split again twice more for the ligation of two more barcodes. After the second ligation, cells were pooled for a third time and split into eight sub-libraries with 12,500 cells/sub-library. Cell counting at this step was done with a manual hemocytometer for accuracy. A fourth barcode was added to each sub-library as the last step of barcoding protocol. Next, the cells were lysed, and barcoded cDNA was amplified, cleaned up, and fragmented. Illumina adapters were added to each sub-library as indexes. A Fragment Analyzer (Agilent) was used to confirm the correct size distribution in the libraries after cDNA amplification and fragmentation steps. 48,000 cells were barcoded for each genotype (H9*_WT_* and H9*_UBE3A m-/p-_*) for the 6 week scRNAseq experiment. 12,000 cells (2,000 cells/biological replicate in each genotype, n = 3) were sequenced at 50,000 reads/cell on an Illumina HiSeq 4000 instrument at 10x configuration (600M reads, 150 PE) for the 6 week experiment (Genewiz). 20,000 cells (2,000 cells/biological replicate in each genotype, n ≥ 3) from the 11 week experiment were submitted to be sequenced at 50,000 reads/cell on a Novaseq 6000 instrument (5B reads, 150 PE) (Genomic Sciences Laboratory, NC State University).

### Data processing for scRNA-seq

Data was processed using an established scRNAseq pipeline (https://combine-lab.github.io/alevin-fry-tutorials/2022/split-seq/). Briefly, raw sequencing data was preprocessed with splitp to pair oligo-dT primer and random hexamer barcodes used in the initial round of barcoding. A splici (spliced + intron) index was created using salmon (v1.7.0) using human GENCODE v39 and hg38 human reference genome. Sequences were mapped to the splici index using salmon and quantified using alevin-fry (v0.5.0). Seurat (v4.0.4) was used to filter, process, and analyze gene expression matrices. The eight sequenced sub-libraries for the 11 week experiment were merged into a single Seurat object and reads not matching sample barcodes within a hamming distance of 1 were filtered out. QC filtering was performed removing cells that did not meet the following criteria: Count_RNA > 2000, nFeature_RNA > 1000, mitochondrial percentage < 10%, resulting in 7,498 high-quality cells for the 6 week experiment and 6,759 high-quality cells for the 11 week experiment for analysis. SCTransform v2 was then used to normalize for read depth across cells, scale the data, and find variable features. The mitochondrial mapping percentage was regressed during normalization to remove this confounding source of variation. Principle component analysis (PCA) was performed and an ElbowPlot was generated to determine the number of informative principal components (50 PCs). FindNeighbors, FindClusters, and UMAP dimensionality reduction visualization were performed. Clusters were annotated based on differentially expressed genes in each cluster as well as known marker genes. Cell types for each cluster were determined using previously identified marker genes. Cell type proportions were calculated for each replicate. Speckle (v1.2.0) was used to compare cell type proportions (n = 3 replicates of 4-5 pooled organoids per genotype, unequal variance) between H9*_WT_* and H9*_UBE3A m-/p-_* organoids using the arcsin square root variance stabilizing transformation propeller test. Please refer to Table S5 for software package information.

### *UBE3A* fractional reads in sequencing data

Raw sequencing data was aligned to hg38 human reference genome using zUMIs (v2.9.4d) processing pipeline. Samtools (v1.10) was then used to calculate the fraction of *UBE3A* reads mapping to exons within the *UBE3A* knockout region (GRCh38/hg38 chr15:25339158-25405461) relative to all *UBE3A* exons (GRCh38/hg38 chr15:25331728-25441024; n = 3 replicates of 4-5 pooled organoids per genotype). Please refer to Table S5 for software package information.

### Organoid size quantification

Images for size experiments were taken using an epifluorescence microscope with a documentation camera with a 4X objective (Nikon Instruments). Quantifications were performed manually in FIJI.

### EdU cell proliferation assay

Assay was performed on EBs and hPSCs using the Click-iT EdU Alexa Fluor 488 Flow Cytometry Assay Kit (Thermo Fisher Scientific). EBs used for the EdU cell proliferation assay were generated and maintained for 5 days using the same protocol as described^31,77^. The manufacturer’s guidelines for the EdU cell proliferation assay were followed. Briefly, EdU was added to the culture media of Day 5 EBs at a final concentration of 10 μM. After 2 hr incubation with EdU, EBs and hPSCs were washed with 1X PBS. Fresh media was then added. After incubating the EBs and hPSCs for 2 hr, they were harvested using Accutase (BioLegend)-mediated dissociation. The cells were washed with 3 mL 1% BSA in PBS and centrifuged at 300 g for 5 min. The cell pellets were resuspended using 100 μL 4% paraformaldehyde in PBS and incubated for 15 min at room temperature, protected from light, followed by 3 mL 1% BSA in PBS wash. The cell pellets were then resuspended in 100 μL 1X Click-iT saponin-based permeabilization and wash reagent and incubated for 15 min at room temperature protected from light. The Click-iT reaction cocktail was prepared per manufacturer guidelines and 500 μL of this cocktail was added to the cells. The reaction mixture was incubated for 30 min at room temperature, protected from light. The cells were washed with 3 mL 1X Click-iT saponin-based permeabilization and wash reagent. The cell pellets were resuspended in 500 μL Click-iT saponin-based permeabilization and wash reagent containing DAPI (Invitrogen). After DAPI staining, samples were loaded into a MACSQuantⓇ VYB flow cytometer (Miltenyi Biotec). FlowJo Software (v10.9.0, Becton Dickenson Life Sciences) was used to analyze flow cytometry data. Please refer to Table S5 for software package information.

### RNA extraction and RT-qPCR

EBs used for measuring *TERT* expression were generated and maintained for 5 days using the same protocol as described^31,77^. hPSCs and Day 5 EBs harvested by Accutase (BioLegend)-mediated dissociation were washed at least 2 times in cold PBS. Total RNA was isolated using Direct-zol RNA MicroPrep Kit (Zymo Research) according to the manufacturer’s protocol. RNA samples were collected in 2 mL RNAse-free tubes and chilled on ice throughout the procedure. cDNA synthesis was performed using 900 ng of total RNA for hPSCs and 1000 ng of total RNA for EBs and the iScript Reverse Transcription Kit (BIO-RAD) according to the manufacturer’s protocol. qPCR reactions were performed using IQ Multiplex Powermix (BIO-RAD) for hPSCs and SsoAdvanced™ Universal SYBR® Green Supermix (BIO-RAD) for EBs on a BIO-RAD 384-well machine (CXF384) with PrimePCR probe assays (BIO-RAD) and primers for TERT (Forward primer: GAAGCCACCTCTTTGGAGGG, Reverse Primer: GAGGAAGTGCTTGGTCTCGG)^80^. Unique assay IDs for UBE3A primers and probe: qHsaCIP0031486. Primer pairs and probes for TBP were custom designed (Table S6). Individual primer pairs and probes were tested before multiplexing reactions. Analysis of UBE3A and TERT expression along with the reference gene TBP was performed in duplicate using Excel by calculating the ΔΔCt value. Data are presented as expression level (2^-ΔΔCt^) relative to TBP.

### Differential gene expression

A Wilcoxon Rank Sum test was used to identify differential gene expression of H9*_UBE3A m-/p-_* cells relative to H9*_WT_* on either all cells or a specific cell subtype based on the analysis. DEG was defined as log2FC≥|1|, and false discovery rate q<0.05.

### Canonical pathway analysis

Ingenuity Pathway Analysis (IPA, version 60467501, QIAGEN Digital Insights) was used to predict activated or inhibited biological functions. Differentially expressed genes with log2(fold change) ≥|1|, false discovery rate q<0.05 were used in IPA. Cancer related biological functions were excluded from the analyses.

### Statistical analysis

All error bars presented represent a 95% confidence interval assuming a normal distribution. p-values for EdU and RT-qPCR experiments were calculated using a with Tukey-Kramer post hoc analysis. p-values for organoid composition and size were calculated using a two tailed t-test. p-value for organoid size was calculated using one tailed t-test. False discovery rates for single cell sequencing were calculated using a Wilcox rank sum test.

## Acknowledgments

We thank Dr. Stormy Chamberlain (University of Connecticut Genetics and Genome Sciences, Farmington, CT) for graciously gifting the H9*_UBE3A m-/p-_* and H9*_UBE3A m-/p+_* cell lines. We thank Dr. Maria Theresa M. Fadri for her thoughtful insights.

## Author contributions

RCE, ZBY, and DS contributed equally to this work. RCE, ZBY, and DS reserve the right to list their name first on professional documents.

Conceptualization: RCE, ZBY, DS, AJK

Methodology: RCE, ZBY, DS, TSP, JMS

Investigation: RCE, ZBY, DS, AJK

Visualization: RCE, ZBY

Supervision: JMS, AJK

Writing—original draft: RCE, ZBY, AJK

Writing—review & editing: RCE, ZBY, DS, TSP, JMS, AJK

## Data and materials availability

Raw data (FASTQ files) and processed data (count matrices) were deposited in NIH GEO (RRID:SCR_005012) under accession no. GSE253230. The code used to perform the analysis described in this study is available at https://github.com/synthetic-keung-lab/scRNAseq_UBE3A_knockout_human_cerebral_organoids. All data needed to evaluate the conclusions in the paper are present in the paper and/or the Supplementary Materials.

## Supplementary Materials

**Fig. S1.**
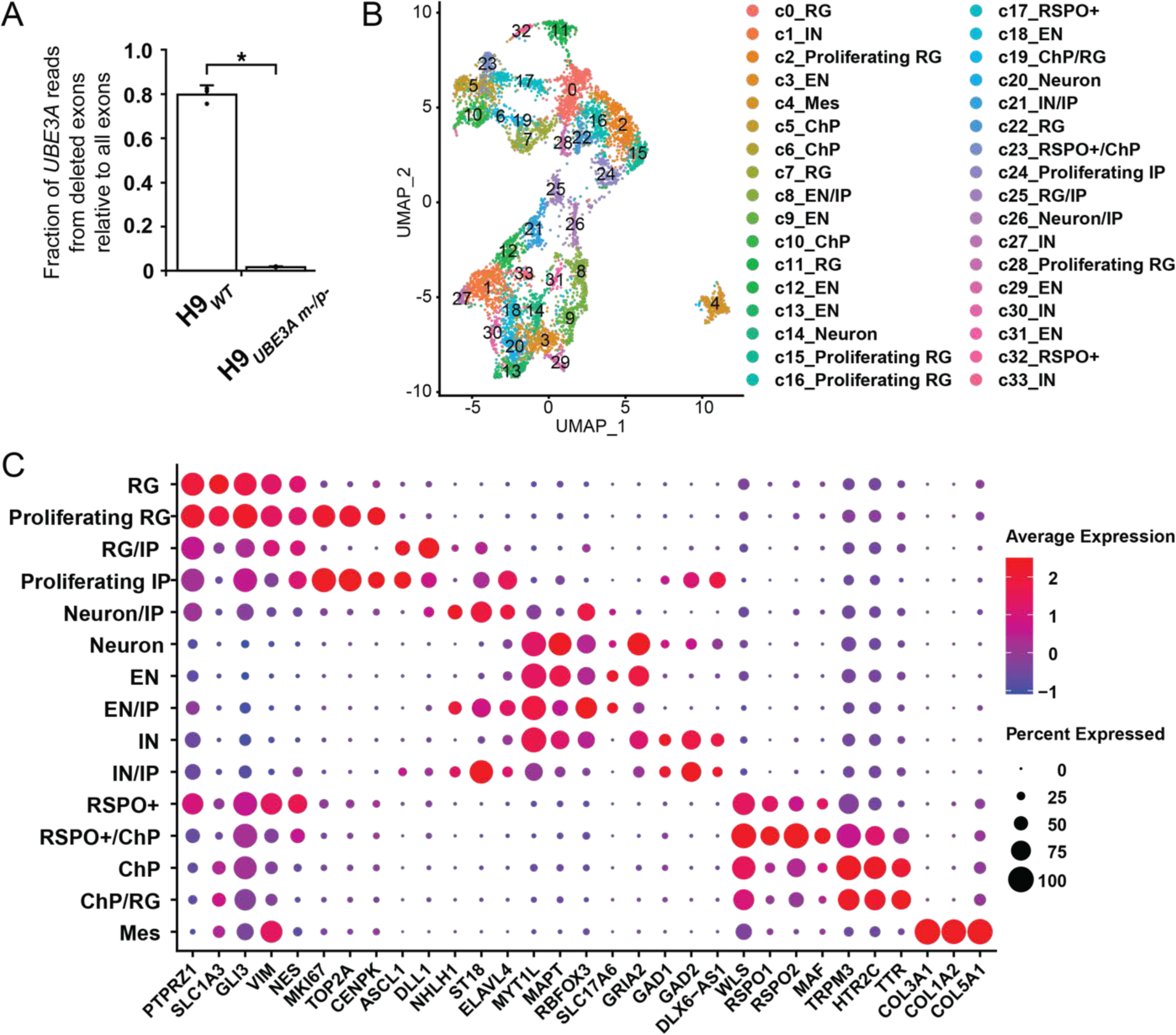
Additional data from the scRNAseq experiment performed on 6 week organoids. Related to Figure 1. (A) Fraction of *UBE3A* reads from deleted exons relative to all exons confirming the deletion in H9*_UBE3A m-/p-_* organoids. Coverage computed for each exon of *UBE3A* for each genotype and replicate for all cells. *p-value <0.05, one-tailed t-test. n = 3 replicates of 4-5 pooled organoids per genotype. Error bars, 95% confidence intervals. (B) UMAP of 6 week H9*_WT_* and H9*_UBE3A m-/p-_* organoid cell cluster identities. RG: radial glia; IN: inhibitory-like neuron; EN: excitatory-like neuron; ChP: choroid plexus; IP: intermediate progenitor; Mes: Mesenchymal. (C) Cell type markers used to identify cell types. There were 15 broad cell types identified consisting of mainly radial glia (RG, e.g., *PTPRZ1*, *GLI3*, *SLC1A3*^81^), intermediate progenitors (IP, e.g., NHLH1^82,83^), mesenchymal cells (e.g., *COL3A1*, *COL1A2*^32^), choroid plexus (ChP, e.g., *TRPM3*, *TTR*^84^), RSPO+ cells (e.g., *RSPO1/2/3*^32^) and neurons (e.g., *MYT1L*, *DCX*^32^). Proliferating RG or IP clusters contained the cell cycle markers (e.g., *MKI67*, *TOP2A*^36^). The neurons containing excitatory (e.g., *SLC17A6*, *GRIA2*^83,85^) and inhibitory (e.g., *GAD1/2*^85,86^) markers were labeled as excitatory-like (EN) and inhibitory-like (IN) neurons, respectively.

**Fig. S2.**
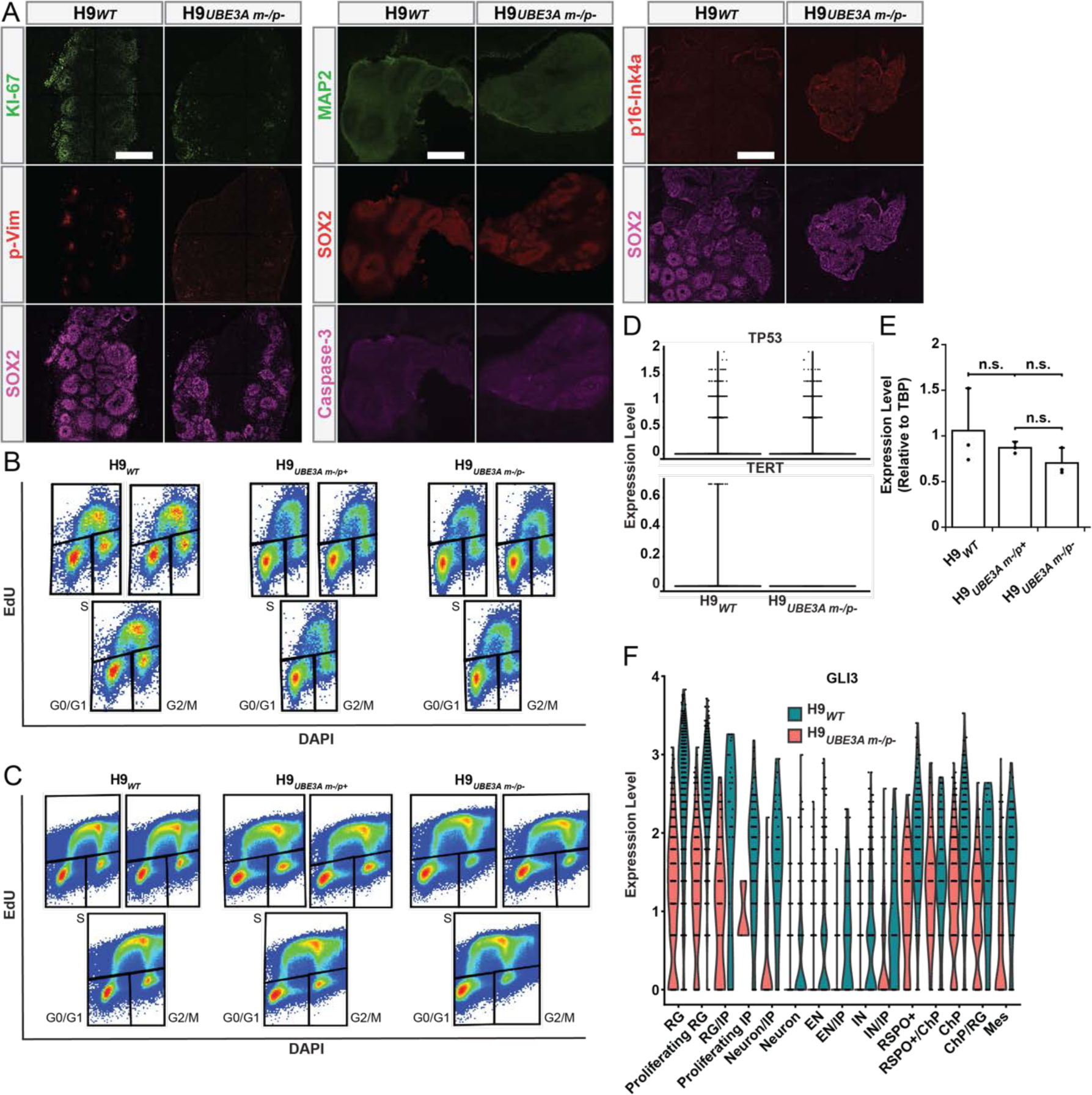
Additional data regarding proliferation, apoptosis, senescence and corticogenesis. Related to Figure 3. (A) Immunostaining of H9_WT_ and H9*_UBE3A m-/p-_* organoids with proliferation (Ki-67, left), apoptosis (Caspase-3, middle) and senescence (p16-Ink4a, right) markers. Scale bars = 500 μm. Flow cytometry plots of EdU labeled H9*_WT_*, H9*_UBE3A m-/p+_* and H9*_UBE3A m-/p-_* (B) embryoid bodies and (C) pluripotent stem cells. (D) Violin plots of *TP53* and *TERT* expression levels for H9*_WT_* and H9*_UBE3A m-/p-_* organoids in scRNAseq data. (E) RT-qPCR measurements of mRNA levels of *TERT* in H9*_WT_*, H9*_UBE3A m-/p+_* and H9*_UBE3A m-/p-_* embryoid bodies normalized to *TBP*, relative to H9*_WT_* embryoid bodies. n.s. (not significant, p-value>0.05) via one-way ANOVA with Tukey-Kramer post hoc analysis, n = 3 replicates of ∼110 pooled embryoid bodies per genotype. Full tick mark indicates sample being compared to half tick mark samples. Error bars are 95% confidence intervals. (F) *GLI3* expression levels in H9*_WT_* and H9*_UBE3A m-/p-_* organoids in scRNAseq data. RG: radial glia; EN: excitatory-like neuron; IN: inhibitory-like neuron; ChP: choroid plexus; IP: intermediate progenitor; EN-CL: cortical layer-like neurons; Mes: mesenchymal; HSP: heat shock protein.

**Fig. S3.**
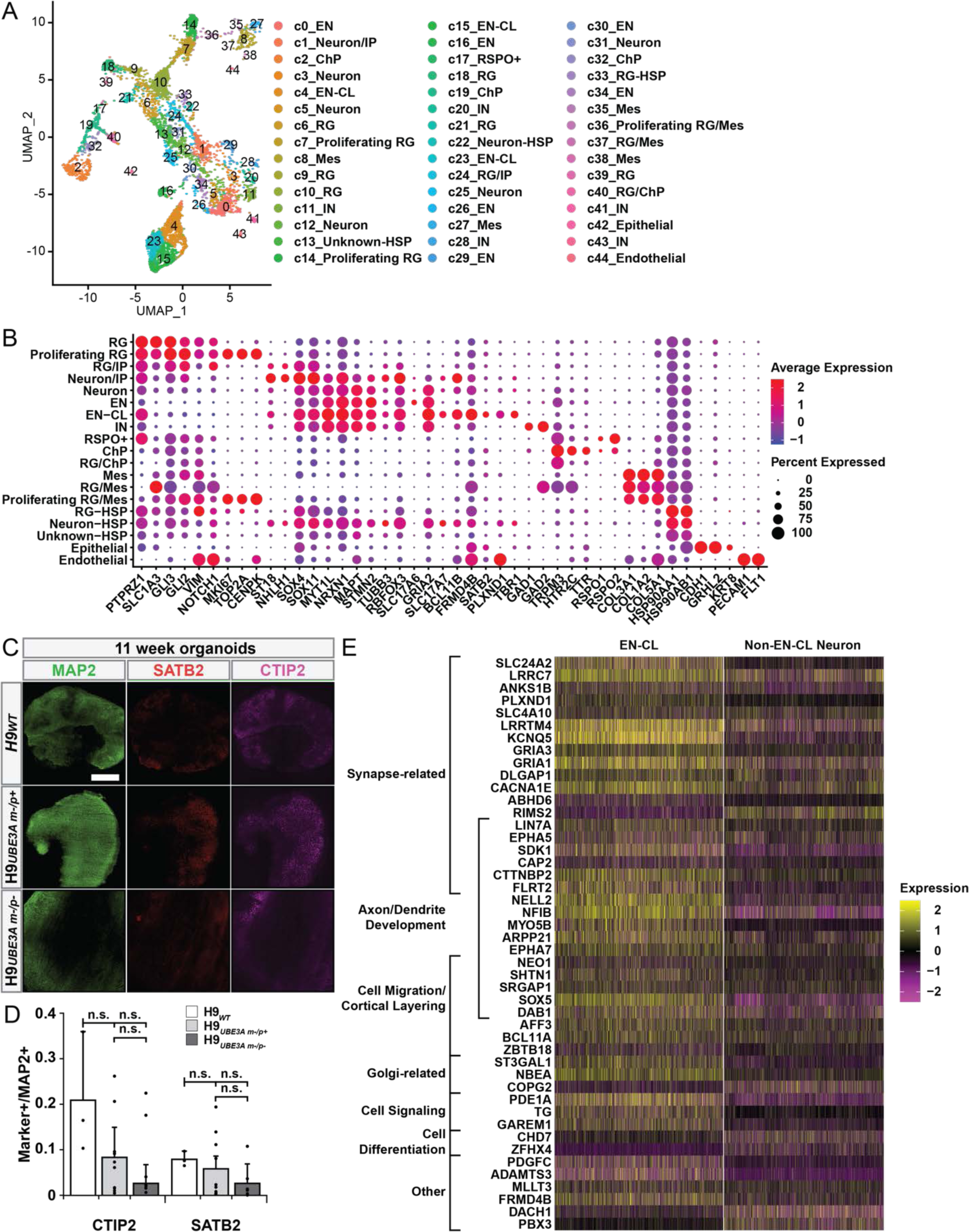
Additional data from the scRNAseq experiment performed on 11 week organoids. Related to Figure 4. (A) UMAP of 11 week H9*_WT_* and H9*_UBE3A m-/p-_* organoid cell cluster identities. RG: radial glia; EN: excitatory-like neuron; IN: inhibitory-like neuron; ChP: choroid plexus; IP: intermediate progenitor; EN-CL: cortical layer-like neurons; Mes: mesenchymal; HSP: heat shock protein. (B) Cell type markers used to identify cell types. There were 18 cell types identified (1 unknown) consisting of mainly radial glia (RG, e.g., *PTPRZ1*, *GLI3*, *SLC1A3*^81^), neuron/intermediate progenitors (neuron/IPs, e.g., *NHLH1*^82,83^), mesenchymal cells (e.g., *COL3A1*, *COL1A2*^32^), choroid plexus (ChP, e.g., *TRPM3*, *TTR*^84^), RSPO+ cells (e.g., *RSPO1/2/3*^32^) and neurons (e.g., *MYT1L*, *DCX*^32^). Proliferating RG or IP clusters contained the cell cycle markers (e.g., *MKI67*, *TOP2A*^36^). Neurons containing excitatory (e.g., *SLC17A6*, *GRIA2*^83,85^) and inhibitory (e.g., *GAD1/2*^85,86^) markers were labeled as excitatory-like (EN) and inhibitory-like (IN) neurons, respectively. Neurons with cortical layer neuron markers (e.g., *FRMD4B*, *SATB2*, *FEZF2*, *PLXND1*, *TBR1*^83,85^) were labeled as EN-CL. Clusters containing heat shock proteins (e.g., *HSP90AA1*, *HSP90AB1*) as top cluster markers were labeled as RG-HSP and Neuron-HSP depending on their marker gene list. There was one cluster unknown with HSPs being the top gene markers (Unknown-HSP). Two small clusters were labeled as epithelial (e.g., *KRT8*^87^, *TJP1*^88^) and endothelial cells (e.g., *FLT1*, *PECAM1*^32^). (C) Immunostaining of EN-CL markers in 11 week H9*_WT_*, H9*_UBE3A m-/p+_* and H9*_UBE3A m-/p-_* organoids. Scale bars = 500 μm. (D) Immunostaining quantification for EN-CL markers in 11 week H9*_WT_*, H9*_UBE3A m-/p+_* and H9*_UBE3A m-/p-_* organoids. n.s. (not significant, p-value>0.05) via one-way ANOVA with Tukey-Kramer post hoc analysis, n ≥ 3 organoids per genotype. Full tick mark indicates sample being compared to half tick mark samples. Error bars are 95% confidence intervals. (E) Heatmap of top differentially expressed markers between EN-CL and non EN-CL neurons (EN, IN, Neuron, Neuron/IP, Neuron-HSP). Differentially expressed markers were grouped by cellular function: synapse-related (*SLC24A2*^89–92^, *LRRC7*^93^, *ANKS1B*^94^, *PLXND1*^95^, *SLC4A10*^96^, *LRRTM4*^97^, *KCNQ5*^98^, *GRIA3*^90–92,99^, *GRIA1*^100^, *DLGAP1*^101^, *CACNA1E*^90–92,102^, *ABHD6*^103,104^, *RIMS2*^105–108^, *LIN7A*^109^, *EPHA5*^110,111^, *SDK1*^90–92,112^, *CAP2*^113,114^, *CTTNBP2*^115,116^, *FLRT2*^90–92,117,118^, *DAB1*^119–121^, *NBEA*^90–92,122,123^), axon/dendrite development (*LIN7A*^109^, *EPHA5*^110,111^, *SDK1*^90–92,112^, *CAP2*^113,114^, *CTTNBP2*^115,116^, *FLRT2*^90–92,117,118^, *NELL2*^124^, *NFIB*^125^, *MYO5B*^126^, *ARPP21*^127^, *EPHA7*^128,129^, *NEO1*^130^, *SHTN1*^131–133^, *SRGAP1*^134^, *SOX5*^135^, *DAB1*^119–121^), cell migration/cortical layering (*LIN7A*^109^, *FLRT2*^90–92,117,118^, *NEO1*^130^, *SHTN1*^131–133^, *SRGAP1*^134^, *SOX5*^135^, *DAB1*^119–121^, *AFF3*^136^, *BCL11A*^137^, *ZBTB18*^138,139^), golgi-related (*ST3GAL1*^90–92,140^, *NBEA*^90–92,122,123^, *COPG2*^141^), cell signaling (*PDE1A*^90–92,142^, *TG*^143^, *GAREM1*^90–92,144,145^), cell differentiation (*CHD7*^146^, *ZFHX4*^147^), and other (*PDGFC*^148,149^, *ADAMTS3*150, *MLLT3*^151^, *FRMD4B*^152^, *DACH1*^153^, *PBX3*^154^).

**Table S1. Qiagen Ingenuity Pathway Analysis canonical pathways for 6 week organoids. Related to Figure 2 (separate file).**

Differentially expressed genes with false discovery rate q < 0.05, log2 (FC) ≥|1| were used in this analysis.

**Table S2. Qiagen Ingenuity Pathway Analysis canonical pathways for 11 week organoids. Related to Figure 5 (separate file).**

Differentially expressed genes with false discovery rate q < 0.05, log2 (FC) ≥|1| were used in this analysis.

**Table S3.**
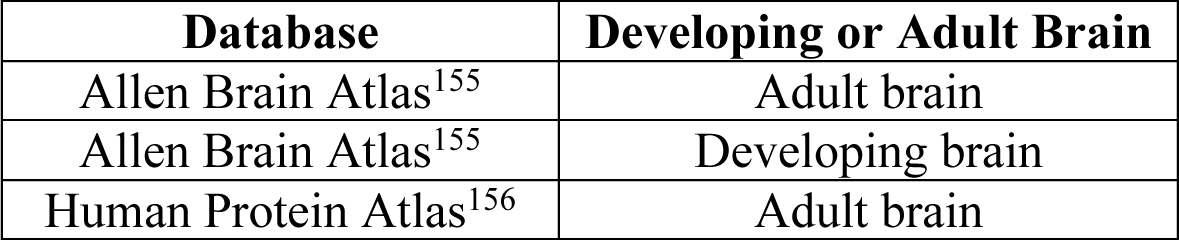
*UBE3A* expression in human ChP tissue database information.

| Database | Developing or Adult Brain |
| --- | --- |
| Allen Brain Atlas<sup>155</sup> | Adult brain |
| Allen Brain Atlas<sup>155</sup> | Developing brain |
| Human Protein Atlas<sup>156</sup> | Adult brain |

**Table S4.** Primary antibodies used in immunostaining experiments.

| Antigen | Host | Supplier | Cat. No. | RRID | Dilution |
| --- | --- | --- | --- | --- | --- |
| SOX2 | Goat | R&D Systems | AF2018 | AB_355110 | 1:20 |
| UBE3A | Rabbit | Bethyl Laboratories | A300-351A | AB_185563 | 1:250 |
| pVimentin | Mouse | MBL International | D076-3S | AB_592962 | 1:200 |
| MAP2 | Mouse | Millipore Sigma | M1406 | AB_477171 | 1:250 |
| MAP2 | Chicken | Abcam | AB92434 | AB_2138147 | 1:300 |
| TTR | Sheep | BIO-RAD | AHP1837 | AB_2212089 | 1:100 |
| EMX1 | Rabbit | Atlas Antibodies | HPA006421 | AB_1078739 | 1:50 |
| Ki-67 | Rabbit | Abcam | AB15580 | AB_443209 | 1:800 |
| Caspase-3 | Rabbit | R&D Systems | AF835 | AB_2243952 | 1:20 |
| p16INK4a | Mouse | Abcam | AB54210 | AB_881819 | 1:1000 |
| CTIP2 | Rat | Abcam | AB18465 | AB_2064130 | 1:100 |
| SATB2 | Mouse | Abcam | AB51502 | AB_882455 | 1:100 |

**Table S5.** Software package information.

| Name | Version |
| --- | --- |
| FlowJo Software<sup>157</sup> | v10.9.0 |
| Salmon<sup>158</sup> | v1.7.0 |
| Alevin-fry<sup>159</sup> | v0.5.0 |
| Seurat<sup>160</sup> | v4.0.4 |
| SCTransform<sup>161</sup> | v2 |
| Speckle<sup>162</sup> | V1.2.0 |
| zUMIs<sup>163</sup> | v2.9.4d |
| Samtools<sup>164</sup> | v1.10 |

**Table S6.**
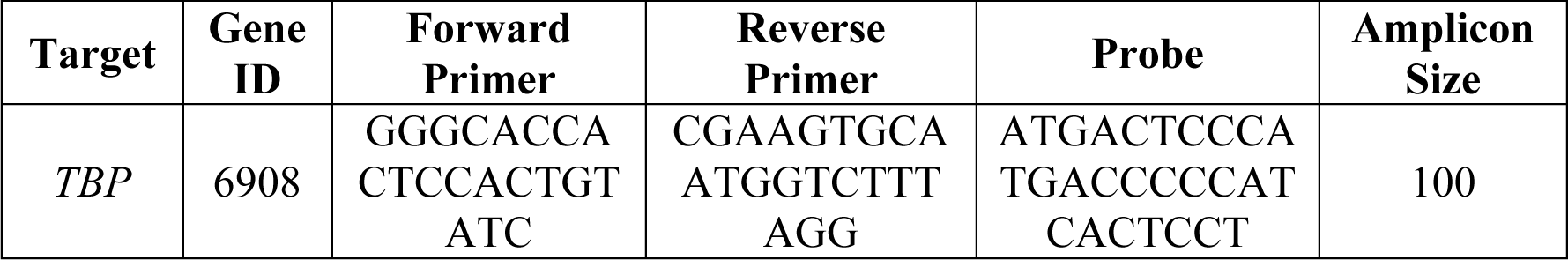
Custom primers used in RT-qPCR experiments.

**Data S1. Raw data related to figures (separate file).**

**Data S2. Raw differential gene expression data related to Figure 2 (separate file).**

**Data S3. Raw differential marker expression data related to Figure 4 (separate file).**

**Data S4. Raw differential gene expression data related to Figure 5 (separate file).**

## References

1. Urraca N, Reiter LT. Developmental Disabilities, Autism, and Schizophrenia at a Single Locus: Complex Gene Regulation and Genomic Instability of 15q11-q13 Cause a Range of Neurodevelopmental Disorders. In: Neural Circuit Development and Function in the Healthy and Diseased Brain: Comprehensive Developmental Neuroscience.; 2013. doi:10.1016/B978-0-12-397267-5.00017-0

2. Rougeulle C, Glatt H, Lalande M. The Angelman syndrome candidate gene, UBE3A/E6-AP, is imprinted in brain. Nat Genet. 1997;17(1). doi:10.1038/ng0997-14

3. Yamasaki K, Joh K, Ohta T, et al. Neurons but not glial cells show reciprocal imprinting of sense and antisense transcripts of Ube3a. Hum Mol Genet. 2003;12(8). doi:10.1093/hmg/ddg106

4. Judson MC, Sosa-Pagan JO, Del Cid WA, Han JE, Philpot BD. Allelic specificity of Ube3a expression in the mouse brain during postnatal Development. J Comp Neurol. 2014;522(8):1874–1896. doi:10.1002/cne.23507

5. Jiang Y hui, Armstrong D, Albrecht U, et al. Mutation of the Angelman ubiquitin ligase in mice causes increased cytoplasmic p53 and deficits of contextual learning and long-term potentiation. Neuron. 1998;21(4). doi:10.1016/S0896-6273(00)80596-6

6. Vatsa N, Jana NR. UBE3A and its link with autism. Front Mol Neurosci. 2018;11. doi:10.3389/fnmol.2018.00448

7. Silva-Santos S, Van Woerden GM, Bruinsma CF, et al. Ube3a reinstatement identifies distinct developmental windows in a murine Angelman syndrome model. J Clin Invest. 2015;125(5). doi:10.1172/JCI80554

8. Wallace ML, Burette AC, Weinberg RJ, Philpot BD. Maternal Loss of Ube3a Produces an Excitatory/Inhibitory Imbalance through Neuron Type-Specific Synaptic Defects. Neuron. 2012;74(5). doi:10.1016/j.neuron.2012.03.036

9. Judson MCC, Wallace MLL, Sidorov MSS, et al. GABAergic Neuron-Specific Loss of Ube3a Causes Angelman Syndrome-Like EEG Abnormalities and Enhances Seizure Susceptibility. Neuron. 2016;90(1). doi:10.1016/j.neuron.2016.02.040

10. Greer PL, Hanayama R, Bloodgood BL, et al. The Angelman Syndrome Protein Ube3A Regulates Synapse Development by Ubiquitinating Arc. Cell. 2010;140(5). doi:10.1016/j.cell.2010.01.026

11. Margolis SS, Salogiannis J, Lipton DM, et al. EphB-mediated degradation of the RhoA GEF Ephexin5 relieves a developmental brake on excitatory synapse formation. Cell. 2010;143(3). doi:10.1016/j.cell.2010.09.038

12. Tonazzini I, Van Woerden GM, Masciullo C, Mientjes EJ, Elgersma Y, Cecchini M. The role of ubiquitin ligase E3A in polarized contact guidance and rescue strategies in UBE3A-deficient hippocampal neurons. Mol Autism. 2019;10(1). doi:10.1186/s13229-019-0293-1

13. Tomaić V, Banks L. Angelman syndrome-associated ubiquitin ligase UBE3A/E6AP mutants interfere with the proteolytic activity of the proteasome. Cell Death Dis. 2015;6(1). doi:10.1038/cddis.2014.572

14. Yi JJ, Berrios J, Newbern JM, et al. An Autism-Linked Mutation Disables Phosphorylation Control of UBE3A. Cell. 2015;162(4). doi:10.1016/j.cell.2015.06.045

15. Yi JJ, Paranjape SR, Walker MP, et al. The autism-linked UBE3A T485A mutant E3 ubiquitin ligase activates the Wnt/β-catenin pathway by inhibiting the proteasome. J Biol Chem. 2017;292(30). doi:10.1074/jbc.M117.788448

16. Xing L, Simon JM, Ptacek TS, et al. Autism-linked UBE3A gain-of-function mutation causes interneuron and behavioral phenotypes when inherited maternally or paternally in mice. Cell Rep. 2023;42(7). doi:10.1016/j.celrep.2023.112706

17. Miao S, Chen R, Ye J, et al. The Angelman Syndrome Protein Ube3a Is Required for Polarized Dendrite Morphogenesis in Pyramidal Neurons. J Neurosci. 2013;33(1):327–333. doi:10.1523/JNEUROSCI.2509-12.2013

18. Dindot S V., Antalffy BA, Bhattacharjee MB, Beaudet AL. The Angelman syndrome ubiquitin ligase localizes to the synapse and nucleus, and maternal deficiency results in abnormal dendritic spine morphology. Hum Mol Genet. 2008;17(1). doi:10.1093/hmg/ddm288

19. Filonova I, Trotter JH, Banko JL, Weeber EJ. Activity-dependent changes in MAPK activation in the Angelman Syndrome mouse model. Learn Mem. 2014;21(2). doi:10.1101/lm.032375.113

20. Judson MC, Burette AC, Thaxton CL, et al. Decreased axon caliber underlies loss of fiber tract integrity, disproportional reductions in white matter volume, and microcephaly in angelman syndrome model mice. J Neurosci. 2017;37(31). doi:10.1523/JNEUROSCI.0037-17.2017

21. Ramamoorthy S, Nawaz Z. E6-associated protein (E6-AP) is a dual function coactivator of steroid hormone receptors. Nucl Recept Signal. 2008;6. doi:10.1621/nrs.06006

22. Nawaz Z, Lonard DM, Smith CL, et al. The Angelman Syndrome-Associated Protein, E6-AP, Is a Coactivator for the Nuclear Hormone Receptor Superfamily. Mol Cell Biol. 1999;19(2). doi:10.1128/mcb.19.2.1182

23. Singhmar P, Kumar A. Angelman syndrome protein UBE3A interacts with primary microcephaly protein ASPM, localizes to centrosomes and regulates chromosome segregation. PLoS One. 2011;6(5). doi:10.1371/journal.pone.0020397

24. Martínez-Noël G, Luck K, Kühnle S, et al. Network Analysis of UBE3A/E6AP-Associated Proteins Provides Connections to Several Distinct Cellular Processes. J Mol Biol. 2018;430(7). doi:10.1016/j.jmb.2018.01.021

25. Herzing LBK, Cook EH, Ledbetter DH. Allele-specific expression analysis by RNA-FISH demonstrates preferential maternal expression of UBE3A and imprint maintenance within 15q11-q13 duplications. Hum Mol Genet. 2002;11(15). doi:10.1093/hmg/11.15.1707

26. Sen D, Voulgaropoulos A, Drobna Z, Keung AJ. Human Cerebral Organoids Reveal Early Spatiotemporal Dynamics and Pharmacological Responses of UBE3A. Stem Cell Reports. Published online 2020. doi:10.1016/j.stemcr.2020.08.006

27. Sonzogni M, Zhai P, Mientjes EJ, Van Woerden GM, Elgersma Y. Assessing the requirements of prenatal UBE3A expression for rescue of behavioral phenotypes in a mouse model for Angelman syndrome. Mol Autism. 2020;11(1). doi:10.1186/s13229-020-00376-9

28. Sonzogni M, Hakonen J, Bernabé Kleijn M, et al. Delayed loss of UBE3A reduces the expression of Angelman syndrome-associated phenotypes. Mol Autism. 2019;10(1). doi:10.1186/s13229-019-0277-1

29. Shen MD, Nordahl CW, Li DD, et al. Extra-axial cerebrospinal fluid in high-risk and normal-risk children with autism aged 2–4 years: a case-control study. The Lancet Psychiatry. 2018;5(11). doi:10.1016/S2215-0366(18)30294-3

30. Dodge A, Willman J, Willman M, et al. Identification of UBE3A Protein in CSF and Extracellular Space of the Hippocampus Suggest a Potential Novel Function in Synaptic Plasticity. Autism Res. 2021;14(4). doi:10.1002/aur.2475

31. Lancaster MA, Renner M, Martin C-A, et al. Cerebral organoids model human brain development and microcephaly. Nature. 2013;501. doi:10.1038/nature12517

32. Camp JG, Badsha F, Florio M, et al. Human cerebral organoids recapitulate gene expression programs of fetal neocortex development. Proc Natl Acad Sci U S A. 2015;112(51):15672–15677. doi:10.1073/pnas.1520760112

33. Logan S, Arzua T, Yan Y, et al. Dynamic Characterization of Structural, Molecular, and Electrophysiological Phenotypes of Human-Induced Pluripotent Stem Cell-Derived Cerebral Organoids, and Comparison with Fetal and Adult Gene Profiles. Cells. 2020;9(5). doi:10.3390/cells9051301

34. Papes F, Camargo AP, de Souza JS, et al. Transcription Factor 4 loss-of-function is associated with deficits in progenitor proliferation and cortical neuron content. Nat Commun. 2022;13(1). doi:10.1038/s41467-022-29942-w

35. Sun AX, Yuan Q, Fukuda M, et al. Potassium channel dysfunction in human neuronal models of Angelman syndrome. Science (80-). 2019;366(6472). doi:10.1126/science.aav5386

36. Kanton S, Boyle MJ, He Z, et al. Organoid single-cell genomic atlas uncovers human-specific features of brain development. Nature. 2019;574(7778). doi:10.1038/s41586-019-1654-9

37. Sirois C. Generation of Isogenic Human Pluripotent Stem Cell-Derived Neurons to Establish a Molecular Angelman Syndrome Phenotype and to Study the UBE3A Protein Isoforms. Published online 2018. https://digitalcommons.lib.uconn.edu/dissertations/1988

38. Azzarelli R, Kerloch T, Pacary E. Regulation of cerebral cortex development by Rho GTPases: insights from in vivo studies. Front Cell Neurosci. 2015;8. https://www.frontiersin.org/articles/10.3389/fncel.2014.00445

39. Denley MCS, Gatford NJF, Sellers KJ, Srivastava DP. Estradiol and the Development of the Cerebral Cortex: An Unexpected Role? Front Neurosci. 2018;12:245. doi:10.3389/fnins.2018.00245

40. McNamara RK. DHA deficiency and prefrontal cortex neuropathology in recurrent affective disorders. J Nutr. 2010;140(4):864–868. doi:10.3945/jn.109.113233

41. Laussu J, Khuong A, Gautrais J, Davy A. Beyond boundaries--Eph:ephrin signaling in neurogenesis. Cell Adh Migr. 2014;8(4):349–359. doi:10.4161/19336918.2014.969990

42. Lin T V, Bordey A. Chapter 15 - mTOR Signaling in Cortical Network Development. In: Faingold CL, Blumenfeld CNS Disorders, and Therapeutics HBT-NN in BF, eds. Academic Press; 2014:193–205. 10.1016/B978-0-12-415804-7.00015-0

43. Frotscher M, Chai X, Bock HH, Haas CA, Förster E, Zhao S. Role of Reelin in the development and maintenance of cortical lamination. J Neural Transm. 2009;116(11):1451–1455. doi:10.1007/s00702-009-0228-7

44. Passemard S, Perez F, Gressens P, El Ghouzzi V. Endoplasmic reticulum and Golgi stress in microcephaly. Cell Stress. 2019;3(12):369–384. doi:10.15698/cst2019.12.206

45. Yeh ML, Gonda Y, Mommersteeg MTM, et al. Robo1 modulates proliferation and neurogenesis in the developing neocortex. J Neurosci Off J Soc Neurosci. 2014;34(16):5717–5731. doi:10.1523/JNEUROSCI.4256-13.2014

46. Liu W, Zhang B, Zhang D, et al. The RBPJ/DAPK3/UBE3A signaling axis induces PBRM1 degradation to modulate the sensitivity of renal cell carcinoma to CDK4/6 inhibitors. Cell Death Dis. 2022;13(4):295. doi:10.1038/s41419-022-04760-6

47. Cheng K-C, Li Y, Chang W-T, Chen Z-C, Cheng J-T, Tsai C-C. Ubiquitin-protein ligase E3a (UBE3A) as a new biomarker of cardiac hypertrophy in cell models. J food drug Anal. 2019;27(1):355–364. doi:10.1016/j.jfda.2018.08.002

48. Simchi L, Panov J, Morsy O, Feuermann Y, Kaphzan H. Novel insights into the role of ube3a in regulating apoptosis and proliferation. J Clin Med. 2020;9(5). doi:10.3390/jcm9051573

49. Mishra A, Godavarthi SK, Jana NR. UBE3A/E6-AP regulates cell proliferation by promoting proteasomal degradation of p27. Neurobiol Dis. 2009;36(1):26–34. doi:10.1016/j.nbd.2009.06.010

50. De Stanchina E, Querido E, Narita M, et al. PML is a direct p53 target that modulates p53 effector functions. Mol Cell. 2004;13(4). doi:10.1016/S1097-2765(04)00062-0

51. Oh W, Ghim J, Lee EW, et al. PML-IV functions as a negative regulator of telomerase by interacting with TERT. J Cell Sci. 2009;122(15). doi:10.1242/jcs.048066

52. Stanurova J, Neureiter A, Hiber M, et al. Angelman syndrome-derived neurons display late onset of paternal UBE3A silencing. Sci Rep. 2016;6. doi:10.1038/srep30792

53. Velasco S, Kedaigle AJ, Simmons SK, et al. Individual brain organoids reproducibly form cell diversity of the human cerebral cortex. Nature. 2019;570(7762):523–527. doi:10.1038/s41586-019-1289-x

54. Chan C-H, Godinho LN, Thomaidou D, Tan S-S, Gulisano M, Parnavelas JG. Emx1 is a Marker for Pyramidal Neurons of the Cerebral Cortex. Cereb Cortex. 2001;11(12):1191–1198. doi:10.1093/cercor/11.12.1191

55. Pollen AA, Nowakowski TJ, Chen J, et al. Molecular identity of human outer radial glia during cortical development. Cell. 2015;163(1):55–67. doi:10.1016/j.cell.2015.09.004

56. Theil T, Alvarez-Bolado G, Walter A, Rüther U. Gli3 is required for Emx gene expression during dorsal telencephalon development. Development. 1999;126(16):3561–3571. doi:10.1242/dev.126.16.3561

57. Ma P, Song N-N, Li Y, et al. Fine-Tuning of Shh/Gli Signaling Gradient by Non-proteolytic Ubiquitination during Neural Patterning. Cell Rep. 2019;28(2):541–553.e4. doi:10.1016/j.celrep.2019.06.017

58. Rougeulle C, Cardoso C, Fontés M, Colleaux L, Lalande M. An imprinted antisense RNA overlaps UBE3A and a second maternally expressed transcript. Nat Genet. 1998;19(1). doi:10.1038/ng0598-15

59. Callen BP, Shearwin KE, Egan JB. Transcriptional interference between convergent promoters caused by elongation over the promoter. Mol Cell. 2004;14(5). doi:10.1016/j.molcel.2004.05.010

60. Crampton N, Bonass WA, Kirkham J, Rivetti C, Thomson NH. Collision events between RNA polymerases in convergent transcription studied by atomic force microscopy. Nucleic Acids Res. 2006;34(19). doi:10.1093/nar/gkl668

61. Osato N, Suzuki Y, Ikeo K, Gojobori T. Transcriptional interferences in cis natural antisense transcripts of humans and mice. Genetics. 2007;176(2). doi:10.1534/genetics.106.069484

62. Pellegrini L, Bonfio C, Chadwick J, Begum F, Skehel M, Lancaster MA. Human CNS barrier-forming organoids with cerebrospinal fluid production. Science (80-). Published online 2020. doi:10.1126/science.aaz5626

63. Estridge RC, O’Neill JE, Keung AJ. Matrigel Tunes H9 Stem Cell-Derived Human Cerebral Organoid Development. Organoids. 2023;2(4):165–176. doi:10.3390/organoids2040013

64. Jacob F, Pather SR, Huang WK, et al. Human Pluripotent Stem Cell-Derived Neural Cells and Brain Organoids Reveal SARS-CoV-2 Neurotropism Predominates in Choroid Plexus Epithelium. Cell Stem Cell. Published online 2020. doi:10.1016/j.stem.2020.09.016

65. Zhang D, Eguchi N, Okazaki S, Sora I, Hishimoto A. Telencephalon Organoids Derived from an Individual with ADHD Show Altered Neurodevelopment of Early Cortical Layer Structure. Stem Cell Rev Reports. 2023;19(5). doi:10.1007/s12015-023-10519-z

66. Simchi L, Gupta PK, Feuermann Y, Kaphzan H. Elevated ROS levels during the early development of Angelman syndrome alter the apoptotic capacity of the developing neural precursor cells. Mol Psychiatry. Published online 2023. doi:10.1038/s41380-023-02038-7

67. Fink JJ, Robinson TM, Germain ND, et al. Disrupted neuronal maturation in Angelman syndrome-derived induced pluripotent stem cells. Nat Commun. 2017;8. doi:10.1038/ncomms15038

68. Yashiro K, Riday TT, Condon KH, et al. Ube3a is required for experience-dependent maturation of the neocortex. Nat Neurosci. 2009;12(6). doi:10.1038/nn.2327

69. Sun J, Liu Y, Jia Y, et al. UBE3A-mediated p18/LAMTOR1 ubiquitination and degradation regulate mTORC1 activity and synaptic plasticity. Elife. 2018;7. doi:10.7554/eLife.37993

70. Yi JJ, Paranjape SR, Walker MP, et al. The autism-linked UBE3A T485A mutant E3 ubiquitin ligase activates the Wnt/β-catenin pathway by inhibiting the proteasome. J Biol Chem. 2017;292(30):12503–12515. doi:10.1074/jbc.M117.788448

71. Vargas T, Ugalde C, Spuch C, et al. Abeta accumulation in choroid plexus is associated with mitochondrial-induced apoptosis. Neurobiol Aging. 2010;31(9):1569–1581. doi:10.1016/j.neurobiolaging.2008.08.017

72. Krzyzanowska A, Carro E. Pathological alteration in the choroid plexus of Alzheimer’s disease: implication for new therapy approaches. Front Pharmacol. 2012;3:75. doi:10.3389/fphar.2012.00075

73. Myung J, Schmal C, Hong S, et al. The choroid plexus is an important circadian clock component. Nat Commun. 2018;9(1):1062. doi:10.1038/s41467-018-03507-2

74. Van Hoecke L, Van Cauwenberghe C, Dominko K, et al. Involvement of the Choroid Plexus in the Pathogenesis of Niemann-Pick Disease Type C. Front Cell Neurosci. 2021;15:757482. doi:10.3389/fncel.2021.757482

75. Nenninger AW, Willman M, Willman J, et al. Improving Gene Therapy for Angelman Syndrome with Secreted Human UBE3A. Neurotherapeutics. 2022;19(4). doi:10.1007/s13311-022-01239-2

76. Singh BK, Vatsa N, Kumar V, Shekhar S, Sharma A, Jana NR. Ube3a deficiency inhibits amyloid plaque formation in APPswe/PS1δE9 mouse model of Alzheimer’s disease. Hum Mol Genet. 2017;26(20):4042–4054. doi:10.1093/hmg/ddx295

77. Lancaster MA, Knoblich JA. Generation of cerebral organoids from human pluripotent stem cells. Nat Protoc. 2014;9(10):2329–2340. doi:10.1038/nprot.2014.158

78. Quadrato G, Nguyen T, Macosko EZ, et al. Cell diversity and network dynamics in photosensitive human brain organoids. Nature. Published online 2017. doi:10.1038/nature22047

79. Rosenberg AB, Roco CM, Muscat RA, et al. Single-cell profiling of the developing mouse brain and spinal cord with split-pool barcoding. Science (80-). 2018;360(6385):176–182. doi:10.1126/science.aam8999

80. Rowland TJ, Dumbović G, Hass EP, Rinn JL, Cech TR. Single-cell imaging reveals unexpected heterogeneity of telomerase reverse transcriptase expression across human cancer cell lines. Proc Natl Acad Sci U S A. 2019;116(37). doi:10.1073/pnas.1908275116

81. Pollen AA, Nowakowski TJ, Chen J, et al. Molecular Identity of Human Outer Radial Glia during Cortical Development. Cell. 2015;163(1). doi:10.1016/j.cell.2015.09.004

82. Jourdon A, Wu F, Mariani J, et al. Modeling idiopathic autism in forebrain organoids reveals an imbalance of excitatory cortical neuron subtypes during early neurogenesis. Nat Neurosci. 2023;26(9):1505–1515. doi:10.1038/s41593-023-01399-0

83. Yao Z, Mich JK, Ku S, et al. A Single-Cell Roadmap of Lineage Bifurcation in Human ESC Models of Embryonic Brain Development. Cell Stem Cell. 2017;20(1). doi:10.1016/j.stem.2016.09.011

84. Diana Neely M, Xie S, Prince LM, et al. Single cell RNA sequencing detects persistent cell type- and methylmercury exposure paradigm-specific effects in a human cortical neurodevelopmental model. Food Chem Toxicol. 2021;154. doi:10.1016/j.fct.2021.112288

85. Trujillo CA, Gao R, Negraes PD, et al. Complex Oscillatory Waves Emerging from Cortical Organoids Model Early Human Brain Network Development. Cell Stem Cell. 2019;25(4):558–569.e7. doi:10.1016/j.stem.2019.08.002

86. van Bruggen D, Pohl F, Langseth CM, et al. Developmental landscape of human forebrain at a single-cell level identifies early waves of oligodendrogenesis. Dev Cell. 2022;57(11). doi:10.1016/j.devcel.2022.04.016

87. Dong J, Hu Y, Fan X, et al. Single-cell RNA-seq analysis unveils a prevalent epithelial/mesenchymal hybrid state during mouse organogenesis. Genome Biol. 2018;19(1). doi:10.1186/s13059-018-1416-2

88. Eze UC, Bhaduri A, Haeussler M, Nowakowski TJ, Kriegstein AR. Single-cell atlas of early human brain development highlights heterogeneity of human neuroepithelial cells and early radial glia. Nat Neurosci. 2021;24(4). doi:10.1038/s41593-020-00794-1

89. Li X-F, Kiedrowski L, Tremblay F, et al. Importance of K^+^-dependent Na^+^/Ca^2+^-exchanger 2, NCKX2, in Motor Learning and Memory *. J Biol Chem. 2006;281(10):6273–6282. doi:10.1074/jbc.M512137200

90. Stelzer G, Rosen N, Plaschkes I, et al. The GeneCards Suite: From Gene Data Mining to Disease Genome Sequence Analyses. Curr Protoc Bioinforma. 2016;54:1.30.1–1.30.33. doi:10.1002/cpbi.5

91. Safran M, Rosen N, Twik M, et al. The GeneCards Suite - Practical Guide to Life Science Databases. In: Abugessaisa I, Kasukawa T, eds. Springer Nature Singapore; 2021:27–56. doi:10.1007/978-981-16-5812-9_2

92. The UniProt Consortium. UniProt: the Universal Protein Knowledgebase in 2023. Nucleic Acids Res. 2023;51(D1):D523–D531. doi:10.1093/nar/gkac1052

93. Izawa I, Nishizawa M, Ohtakara K, Inagaki M. Densin-180 Interacts with &#x3b4;-Catenin/Neural Plakophilin-related Armadillo Repeat Protein at Synapses *. J Biol Chem. 2002;277(7):5345–5350. doi:10.1074/jbc.M110052200

94. Cho CH, Deyneko IV, Cordova-Martinez D, et al. ANKS1B encoded AIDA-1 regulates social behaviors by controlling oligodendrocyte function. Nat Commun. 2023;14(1):8499. doi:10.1038/s41467-023-43438-1

95. Wang F, Eagleson KL, Levitt P. Positive regulation of neocortical synapse formation by the Plexin-D1 receptor. Brain Res. 2015;1616:157–165. doi:10.1016/j.brainres.2015.05.005

96. Sinning A, Liebmann L, Hübner CA. Disruption of Slc4a10 augments neuronal excitability and modulates synaptic short-term plasticity. Front Cell Neurosci. 2015;9:223. doi:10.3389/fncel.2015.00223

97. Sinha R, Siddiqui TJ, Padmanabhan N, et al. LRRTM4: A Novel Regulator of Presynaptic Inhibition and Ribbon Synapse Arrangements of Retinal Bipolar Cells. Neuron. 2020;105(6):1007–1017.e5. doi:10.1016/j.neuron.2019.12.028

98. Schroeder BC, Hechenberger M, Weinreich F, Kubisch C, Jentsch TJ. KCNQ5, a Novel Potassium Channel Broadly Expressed in Brain, Mediates M-type Currents*. J Biol Chem. 2000;275(31):24089–24095. 10.1074/jbc.M003245200

99. Wu Y, Arai AC, Rumbaugh G, et al. Mutations in ionotropic AMPA receptor 3 alter channel properties and are associated with moderate cognitive impairment in humans. Proc Natl Acad Sci. 2007;104(46):18163–18168. doi:10.1073/pnas.0708699104

100. Andrásfalvy BK, Smith MA, Borchardt T, Sprengel R, Magee JC. Impaired regulation of synaptic strength in hippocampal neurons from GluR1-deficient mice. J Physiol. 2003;552(Pt 1):35–45. doi:10.1113/jphysiol.2003.045575

101. Rasmussen AH, Rasmussen HB, Silahtaroglu A. The DLGAP family: neuronal expression, function and role in brain disorders. Mol Brain. 2017;10(1):43. doi:10.1186/s13041-017-0324-9

102. Helbig KL, Lauerer RJ, Bahr JC, et al. De Novo Pathogenic Variants in CACNA1E Cause Developmental and Epileptic Encephalopathy with Contractures, Macrocephaly, and Dyskinesias. Am J Hum Genet. 2018;103(5):666–678. doi:10.1016/j.ajhg.2018.09.006

103. Cao JK, Kaplan J, Stella N. ABHD6: Its Place in Endocannabinoid Signaling and Beyond. Trends Pharmacol Sci. 2019;40(4):267–277. doi:10.1016/j.tips.2019.02.002

104. Wei M, Zhang J, Jia M, et al. α/β-Hydrolase domain-containing 6 (ABHD6) negatively regulates the surface delivery and synaptic function of AMPA receptors. Proc Natl Acad Sci U S A. 2016;113(19):E2695–704. doi:10.1073/pnas.1524589113

105. Mechaussier S, Almoallem B, Zeitz C, et al. Loss of Function of RIMS2 Causes a Syndromic Congenital Cone-Rod Synaptic Disease with Neurodevelopmental and Pancreatic Involvement. Am J Hum Genet. 2020;106(6):859–871. doi:10.1016/j.ajhg.2020.04.018

106. Acuna C, Liu X, Südhof TC. How to Make an Active Zone: Unexpected Universal Functional Redundancy between RIMs and RIM-BPs. Neuron. 2016;91(4):792–807. doi:10.1016/j.neuron.2016.07.042

107. Gebhart M, Juhasz-Vedres G, Zuccotti A, et al. Modulation of Cav1.3 Ca2+ channel gating by Rab3 interacting molecule. Mol Cell Neurosci. 2010;44(3):246–259. doi:10.1016/j.mcn.2010.03.011

108. Li X, Cai D, Huang Y, et al. Aberrant methylation in neurofunctional gene serves as a hallmark of tumorigenesis and progression in colorectal cancer. BMC Cancer. 2023;23(1):315. doi:10.1186/s12885-023-10765-x

109. Matsumoto A, Mizuno M, Hamada N, et al. LIN7A depletion disrupts cerebral cortex development, contributing to intellectual disability in 12q21-deletion syndrome. PLoS One. 2014;9(3):e92695. doi:10.1371/journal.pone.0092695

110. Cooper MA, Crockett DP, Nowakowski RS, Gale NW, Zhou R. Distribution of EphA5 receptor protein in the developing and adult mouse nervous system. J Comp Neurol. 2009;514(4):310–328. doi:10.1002/cne.22030

111. Akaneya Y, Sohya K, Kitamura A, et al. Ephrin-A5 and EphA5 interaction induces synaptogenesis during early hippocampal development. PLoS One. 2010;5(8):e12486. doi:10.1371/journal.pone.0012486

112. Rochon P-L, Theriault C, Rangel Olguin AG, Krishnaswamy A. The cell adhesion molecule Sdk1 shapes assembly of a retinal circuit that detects localized edges. Elife. 2021;10. doi:10.7554/eLife.70870

113. Kumar A, Paeger L, Kosmas K, Kloppenburg P, Noegel AA, Peche VS. Neuronal Actin Dynamics, Spine Density and Neuronal Dendritic Complexity Are Regulated by CAP2. Front Cell Neurosci. 2016;10. https://www.frontiersin.org/articles/10.3389/fncel.2016.00180

114. Pelucchi S, Vandermeulen L, Pizzamiglio L, et al. Cyclase-associated protein 2 dimerization regulates cofilin in synaptic plasticity and Alzheimer’s disease. Brain Commun. 2020;2(2):fcaa086. doi:10.1093/braincomms/fcaa086

115. Chen Y-K, Chen C-Y, Hu H-T, Hsueh Y-P. CTTNBP2, but not CTTNBP2NL, regulates dendritic spinogenesis and synaptic distribution of the striatin-PP2A complex. Mol Biol Cell. 2012;23(22):4383–4392. doi:10.1091/mbc.E12-05-0365

116. Shih P-Y, Hsieh B-Y, Lin M-H, et al. CTTNBP2 Controls Synaptic Expression of Zinc-Related Autism-Associated Proteins and Regulates Synapse Formation and Autism-like Behaviors. Cell Rep. 2020;31(9):107700. doi:10.1016/j.celrep.2020.107700

117. Yamagishi S, Hampel F, Hata K, et al. FLRT2 and FLRT3 act as repulsive guidance cues for Unc5-positive neurons. EMBO J. 2011;30(14):2920–2933. doi:10.1038/emboj.2011.189

118. Cicvaric A, Yang J, Bulat T, et al. Enhanced synaptic plasticity and spatial memory in female but not male FLRT2-haplodeficient mice. Sci Rep. 2018;8(1):3703. doi:10.1038/s41598-018-22030-4

119. O’Dell RS, Ustine CJM, Cameron DA, et al. Layer 6 cortical neurons require Reelin-Dab1 signaling for cellular orientation, Golgi deployment, and directed neurite growth into the marginal zone. Neural Dev. 2012;7(1):25. doi:10.1186/1749-8104-7-25

120. Hammond V, Howell B, Godinho L, Tan SS. disabled-1 functions cell autonomously during radial migration and cortical layering of pyramidal neurons. J Neurosci Off J Soc Neurosci. 2001;21(22):8798–8808. doi:10.1523/JNEUROSCI.21-22-08798.2001

121. Trotter J, Lee GH, Kazdoba TM, et al. Dab1 is required for synaptic plasticity and associative learning. J Neurosci Off J Soc Neurosci. 2013;33(39):15652–15668. doi:10.1523/JNEUROSCI.2010-13.2013

122. Miller AC, Voelker LH, Shah AN, Moens CB. Neurobeachin is required postsynaptically for electrical and chemical synapse formation. Curr Biol. 2015;25(1):16–28. doi:10.1016/j.cub.2014.10.071

123. Nair R, Lauks J, Jung S, et al. Neurobeachin regulates neurotransmitter receptor trafficking to synapses. J Cell Biol. 2013;200(1):61–80. doi:10.1083/jcb.201207113

124. Kim HR, Kim DH, An JY, et al. NELL2 Function in Axon Development of Hippocampal Neurons. Mol Cells. 2020;43(6):581–589. doi:10.14348/molcells.2020.0032

125. Betancourt J, Katzman S, Chen B. Nuclear factor one B regulates neural stem cell differentiation and axonal projection of corticofugal neurons. J Comp Neurol. 2014;522(1):6–35. doi:10.1002/cne.23373

126. Liu Y, Xu X-H, Chen Q, et al. Myosin Vb controls biogenesis of post-Golgi Rab10 carriers during axon development. Nat Commun. 2013;4(1):2005. doi:10.1038/ncomms3005

127. Rehfeld F, Maticzka D, Grosser S, et al. The RNA-binding protein ARPP21 controls dendritic branching by functionally opposing the miRNA it hosts. Nat Commun. 2018;9(1):1235. doi:10.1038/s41467-018-03681-3

128. Leonard CE, Baydyuk M, Stepler MA, Burton DA, Donoghue MJ. EphA7 isoforms differentially regulate cortical dendrite development. PLoS One. 2020;15(12):e0231561. doi:10.1371/journal.pone.0231561

129. Clifford MA, Athar W, Leonard CE, et al. EphA7 signaling guides cortical dendritic development and spine maturation. Proc Natl Acad Sci. 2014;111(13):4994–4999.doi:10.1073/pnas.1323793111

130. Villanueva AA, Falcón P, Espinoza N, et al. The Netrin-4/Neogenin-1 axis promotes neuroblastoma cell survival and migration. Oncotarget. 2017;8(6):9767–9782. doi:10.18632/oncotarget.14213

131. Emerson SE, Stergas HR, Bupp-Chickering SO, Ebert AM. Shootin-1 is required for nervous system development in zebrafish. Dev Dyn an Off Publ Am Assoc Anat. 2020;249(10):1285–1295. doi:10.1002/dvdy.194

132. Zhang M, Ergin V, Lin L, Stork C, Chen L, Zheng S. Axonogenesis Is Coordinated by Neuron-Specific Alternative Splicing Programming and Splicing Regulator PTBP2. Neuron. 2019;101(4):690–706.e10. doi:10.1016/j.neuron.2019.01.022

133. Ergin V, Erdogan M, Menevse A. Regulation of Shootin1 Gene Expression Involves NGF-induced Alternative Splicing during Neuronal Differentiation of PC12 Cells. Sci Rep. 2015;5(1):17931. doi:10.1038/srep17931

134. Lucas B, Hardin J. Mind the (sr)GAP - roles of Slit-Robo GAPs in neurons, brains and beyond. J Cell Sci. 2017;130(23):3965–3974. doi:10.1242/jcs.207456

135. Kwan KY, Lam MMS, Krsnik Z, Kawasawa YI, Lefebvre V, Sestan N. SOX5 postmitotically regulates migration, postmigratory differentiation, and projections of subplate and deep-layer neocortical neurons. Proc Natl Acad Sci U S A. 2008;105(41):16021–16026. doi:10.1073/pnas.0806791105

136. Moore JM, Oliver PL, Finelli MJ, et al. Laf4/Aff3, a gene involved in intellectual disability, is required for cellular migration in the mouse cerebral cortex. PLoS One. 2014;9(8):e105933. doi:10.1371/journal.pone.0105933

137. Wiegreffe C, Simon R, Peschkes K, et al. Bcl11a (Ctip1) Controls Migration of Cortical Projection Neurons through Regulation of Sema3c. Neuron. 2015;87(2):311–325. doi:10.1016/j.neuron.2015.06.023

138. Cohen JS, Srivastava S, Farwell Hagman KD, et al. Further evidence that de novo missense and truncating variants in ZBTB18 cause intellectual disability with variable features. Clin Genet. 2017;91(5):697–707. 10.1111/cge.12861

139. Wang R, Bhatt AB, Minden-Birkenmaier BA, et al. ZBTB18 restricts chromatin accessibility and prevents transcriptional adaptations that drive metastasis. Sci Adv. 2024;9(1):eabq3951. doi:10.1126/sciadv.abq3951

140. Whitehouse C, Burchell J, Gschmeissner S, Brockhausen I, Lloyd KO, Taylor-Papadimitriou J. A transfected sialyltransferase that is elevated in breast cancer and localizes to the medial/trans-Golgi apparatus inhibits the development of core-2-based O-glycans. J Cell Biol. 1997;137(6):1229–1241. doi:10.1083/jcb.137.6.1229

141. Steiner A, Hrovat-Schaale K, Prigione I, et al. Deficiency in coatomer complex I causes aberrant activation of STING signalling. Nat Commun. 2022;13(1):2321. doi:10.1038/s41467-022-29946-6

142. Michibata H, Yanaka N, Kanoh Y, Okumura K, Omori K. Human Ca2+/calmodulin-dependent phosphodiesterase PDE1A: novel splice variants, their specific expression, genomic organization, and chromosomal localization. Biochim Biophys Acta - Gene Struct Expr. 2001;1517(2):278–287. 10.1016/S0167-4781(00)00293-1

143. Sue M, Hayashi M, Kawashima A, et al. Thyroglobulin (Tg) activates MAPK pathway to induce thyroid cell growth in the absence of TSH, insulin and serum. Biochem Biophys Res Commun. 2012;420(3):611–615. doi:10.1016/j.bbrc.2012.03.046

144. Nishino T, Abe T, Kaneko M, et al. GAREM1 is involved in controlling body mass in mice and humans. Biochem Biophys Res Commun. 2022;628:91–97. 10.1016/j.bbrc.2022.08.058

145. Tashiro K, Tsunematsu T, Okubo H, et al. GAREM, a Novel Adaptor Protein for Growth Factor Receptor-bound Protein 2, Contributes to Cellular Transformation through the Activation of Extracellular Signal-regulated Kinase Signaling*. J Biol Chem. 2009;284(30):20206–20214. 10.1074/jbc.M109.021139

146. Yao H, Hannum DF, Zhai Y, et al. CHD7 promotes neural progenitor differentiation in embryonic stem cells via altered chromatin accessibility and nascent gene expression. Sci Rep. 2020;10(1):17445. doi:10.1038/s41598-020-74537-4

147. Hemmi K, Ma D, Miura Y, et al. A homeodomain-zinc finger protein, ZFHX4, is expressed in neuronal differentiation manner and suppressed in muscle differentiation manner. Biol Pharm Bull. 2006;29(9):1830–1835. doi:10.1248/bpb.29.1830

148. Hamada T, Ui-Tei K, Imaki J, et al. The expression of SCDGF/PDGF-C/fallotein and SCDGF-B/PDGF-D in the rat central nervous system. Mech Dev. 2002;112(1-2):161–164. doi:10.1016/s0925-4773(01)00625-6

149. Tian Y, Zhan Y, Jiang Q, Lu W, Li X. Expression and function of PDGF-C in development and stem cells. Open Biol. 2021;11(12):210268. doi:10.1098/rsob.210268

150. Ogino H, Hisanaga A, Kohno T, et al. Secreted Metalloproteinase ADAMTS-3 Inactivates Reelin. J Neurosci Off J Soc Neurosci. 2017;37(12):3181–3191. doi:10.1523/JNEUROSCI.3632-16.2017

151. Büttner N, Johnsen SA, Kügler S, Vogel T. Af9/Mllt3 interferes with Tbr1 expression through epigenetic modification of histone H3K79 during development of the cerebral cortex. Proc Natl Acad Sci U S A. 2010;107(15):7042–7047. doi:10.1073/pnas.0912041107

152. Kumamoto T, Hanashima C. Neuronal subtype specification in establishing mammalian neocortical circuits. Neurosci Res. 2014;86:37–49. doi:10.1016/j.neures.2014.07.002

153. Castiglioni V, Faedo A, Onorati M, et al. Dynamic and Cell-Specific DACH1 Expression in Human Neocortical and Striatal Development. Cereb Cortex. 2019;29(5):2115–2124. doi:10.1093/cercor/bhy092

154. Rottkamp CA, Lobur KJ, Wladyka CL, Lucky AK, O’Gorman S. Pbx3 is required for normal locomotion and dorsal horn development. Dev Biol. 2008;314(1):23–39. doi:10.1016/j.ydbio.2007.10.046

155. Sunkin SM, Ng L, Lau C, et al. Allen Brain Atlas: An integrated spatio-temporal portal for exploring the central nervous system. Nucleic Acids Res. 2013;41(D1). doi:10.1093/nar/gks1042

156. Pontén F, Jirström K, Uhlen M. The Human Protein Atlas - A tool for pathology. J Pathol. 2008;216(4). doi:10.1002/path.2440

157. FlowJo^TM^ Software (for Windows), v10.9.0. Ashland, OR: Becton, Dickinson and Company; 2023.

158. Patro R, Duggal G, Love MI, Irizarry RA, Kingsford C. Salmon: fast and bias-aware quantification of transcript expression using dual-phase inference. Nat Methods. 2017;14(4).

159. He D, Zakeri M, Sarkar H, Soneson C, Srivastava A, Patro R. Alevin-fry unlocks rapid, accurate and memory-frugal quantification of single-cell RNA-seq data. Nat Methods. 2022;19(3). doi:10.1038/s41592-022-01408-3

160. Butler A, Hoffman P, Smibert P, Papalexi E, Satija R. Integrating single-cell transcriptomic data across different conditions, technologies, and species. Nat Biotechnol. 2018;36(5):411–420. doi:10.1038/nbt.4096

161. Hafemeister C, Satija R. Normalization and variance stabilization of single-cell RNA-seq data using regularized negative binomial regression. Genome Biol. 2019;20(1). doi:10.1186/s13059-019-1874-1

162. Phipson B, Sim CB, Porrello ER, Hewitt AW, Powell J, Oshlack A. propeller: testing for differences in cell type proportions in single cell data. Bioinformatics. 2022;38(20). doi:10.1093/bioinformatics/btac582

163. Parekh S, Ziegenhain C, Vieth B, Enard W, Hellmann I. zUMIs - A fast and flexible pipeline to process RNA sequencing data with UMIs. Gigascience. 2018;7(6):1–9. doi:10.1093/gigascience/giy059

164. Li H, Handsaker B, Wysoker A, et al. The Sequence Alignment/Map format and SAMtools. Bioinformatics. 2009;25(16). doi:10.1093/bioinformatics/btp352

